# A lipid plug affects K_2P_6.1(TWIK-2) function

**DOI:** 10.1101/2025.06.11.659167

**Authors:** Abhisek Mondal, Sangeeta Niranjan, Daniel L. Minor

## Abstract

Lipids are integral to ion channel function yet delineating mechanisms by which they affect function remains challenging. Within the K_2P_ family of leak potassium channels^1–3^, observation of tubular densities interpreted as alkyl chains occupying lateral fenestrations linking the pore and bilayer^4–8^ raised the possibility that lipid access from the bilayer acts as a regulatory mechanism^4–7^. Here, we present cryo-electron microscopy (cryo-EM) structures of the human leak potassium channel K_2P_6.1 (TWIK2)^9–11^ and mutants in nanodisc and detergent environments that reveal an unusual conformation in the first selectivity filter (SF1) and a pair of two-chain lipids within the channel cavity (denoted the ‘lipid plug’). The chains of each plug lipid occupy separate binding sites that laterally extend to the bilayer from the channel cavity. One, the upper leg, matches the previously identified alkyl chain binding site^4–8,12^. The second, the lower leg, occupies a fenestration common with K_2P_1.1 (TWIK1)^13^. Together, they demonstrate a bidentate means to coordinate each plug lipid that offers a reinterpretation of previous observations. Structures of a K_2P_6.1 (TWIK2) mutant that directs the channel to the plasma membrane^14^ and an R257A mutant that increases function yield plugged and unplugged forms. Notably, the R257A plugged form shows a change in lipid plug position, indicating a key role for this residue in lipid binding. Together, our data suggest that occupation of the central cavity by the lipid plug serves as a mechanism to render the TWIK channels inactive and points to the importance of lipid plug removal to create an ion permeable pore. Such a mechanism could provide a potent way for limiting the leak function of K_2P_s based on cellular location or other contextual factors.

## Introduction

K_2P_ potassium channels produce leak currents that are important for controlling resting membrane potential in diverse cell types^1–3^. Fifteen human K_2P_ channel subunits comprise six subfamilies (TWIK, TREK, THIK, TASK, TALK, and TRESK) that each respond to diverse sets of physiological cues including physical force, lipid modulation, and signals from various signaling cascades^15^. Structural studies of members from the TWIK^4,5^, TREK^8,16,17^, TALK^18^, TASK^19^, and THIK^20,21^ subfamilies reveal a shared architecture^15^ in which the two pore domains (PD1 and PD2) of an individual K_2P_s subunit form a pseudo tetrameric pore upon subunit dimerization. In contrast to other types of potassium channels, the selectivity filter (SF) ‘C-type’ gate acts as the principal site of K_2P_ modulation for most K_2P_s^15,16,22–26^. Despite their relatively small size (∼60-70 kDa), K_2P_s contain a wealth of binding sites for various classes of lipid and small molecule modulators whose roles in channel gating remain imperfectly understood^15^. In particular, the potential role of lipid access through a lateral fenestration below the selectivity filter has attracted much attention^5,6^, but whether this represents a mechanism for direct block of the pore remains a point of controversy as simulations indicate that even though lipid tails can enter they do not block the pore^13,27^.

K_2P_6.1 (TWIK2)^9–11^ belongs to the TWIK subfamily comprising K_2P_1.1 (TWIK1), K_2P_6.1 (TWIK2), and K_2P_7.1^2^ and is found in the gastrointestinal tract, vasculature, and immune system^9^ where it is linked to vascular^28^ and pulmonary^29^ hypertension and sepsis responses^30^. K_2P_6.1 (TWIK2) and K_2P_1.1 (TWIK1) are notable for their endolysomal localization^31^. Functional and cell biology studies indicate that in K_2P_6.1 (TWIK2) this intracellular localization originates from internalization signals that favor this location and largely prevent plasma membrane expression^14^. Interestingly, in macrophages transit of endosomally sequestered K_2P_6.1 (TWIK2) to the plasma membrane following a signal initiated by extracellular ATP has been implicated in the process underlying NLRP3 (<u>N</u>OD, <u>L</u>RR and <u>p</u>yrin domain-containing protein <u>3</u>) inflammasome activation^30,32–34^. This K_2P_6.1 (TWIK2) function parallels that described for the THIK subfamily in NLRP3 activation in microglia and interleukin 1β (IL–1β) release^35–37^, highlighting the importance of the link between K_2P_ cellular location and function. Although K_2P_1.1 (TWIK-1) has been structurally characterized^4,5,18^, the structure of K_2P_6.1 (TWIK2) has not been previously described.

Here, we report structural studies of human K_2P_6.1 (TWIK2) and mutants in lipid nanodisc and detergent environments. The data reveal a channel architecture that largely resembles K_2P_1.1 (TWIK1) having three notable features: an extra helix in the extracellular cap domain (the ‘ear helix’), an unusual conformation of a key residue in the first selectivity filter (SF1), and block by pair of intracellular lipids that engage the channel through a set of bidentate interactions with lateral fenestrations linking the pore and bilayer. Elements from this last feature occupy the site that has been identified in K_2P_1.1 (TWIK1) ^4,5^ and other K_2P_s ^6–8^ as binding site for a single alkyl chain^4,6–8^ or detergent^4,12^. Structures of a previously characterized K_2P_6.1 (TWIK2), a mutant that increases the plasma membrane localization and activity^14^, yield plugged and unplugged forms that highlight the importance of lipid removal for channel function. Moreover, structure of the R257A mutant that enhances function and affects a key lipid coordinating residue reveals a descent of the lipid plug towards the intracellular opening, highlighting the important role of Arg257 in lipid coordination. The fact that the lipid plug forms a physical barrier that prevents ion passage and that this plug can be destabilized by mutations that enhance channel function suggests that the function of the TWIK subfamily depends on lipid plug removal.

## Results

### Human K_2P_6.1(TWIK-2) structure reveals an unusual selectivity filter conformation and a lipid plug

Expression and purification of full-length human K_2P_6.1(TWIK-2) (denoted ‘TWIK2’) from HEK293 cells yielded a sample suitable for cryo-EM studies (Fig. S1a) that we used for structural characterization in MSP1E1^38^ lipid nanodiscs under high potassium (200 mM KCl) conditions (Figs. S1 and S2, Table 1). The structure (TWIK2:ND) has an overall resolution of 3.2Å, with the best-defined parts reaching 2.2Å (Fig. 1a and S2c, Table 1). TWIK2 has the canonical architecture found throughout the K_2P_ family^15^ (Fig. S3) and is very similar to its related TWIK subfamily member, K_2P_1.1 (TWIK-1)^4,5^ (RMSD_Cα_ = 0.817Å 3UKM^4^, 1.836Å 7SK0^5^, 1.338Å 7SK1^5^). The two PDs from each subunit are arranged around the central axis of the channel with the M2 and M4 helices lining the pore and the M1 and M3 helices located on the periphery. The M1 helix is domain swapped between the two subunits and connects to the extracellular Cap domain as in other K_2P_s ^15^. There is a branched aqueous pathway, the extracellular ion pathway (EIP), between the Cap and extracellular face of the pore that connects the external mouth of the channel selectivity filter (SF) with the extracellular space^4,7,15^. Different from other K_2P_s, the TWIK2 Cap domain has a third helix (denoted the ‘ear helix’) that connects the Cap to the loop leading to the P1 helix (Fig. 1b, S4a-b). Ear helix residues Gly73, Val76, Leu77, and Asn79 from one subunit pack against the α2 helix from its own Cap and the α1 and α2 Cap helices from its dimeric partner. These positions are largely conserved among vertebrate TWIK2s (Fig. S5) and with exception of a hydrogen bond pair (Glu41-Asn79) are conserved in TWIK1 (Fig. S4b). By contrast, these sites are poorly conserved in other K_2P_ classes (Figs. S4a-b), indicating that the Ear helix is a feature of the TWIK subfamily. Evaluation of surface electrostatics highlights the electronegative character of the extracellular EIP, a prominent positively charged surface on the intracellular M2-M3 loop, and a relatively hydrophobic inner cavity bounded by the selectivity filter and Arg257 (Fig. S6).

**Figure 1.**
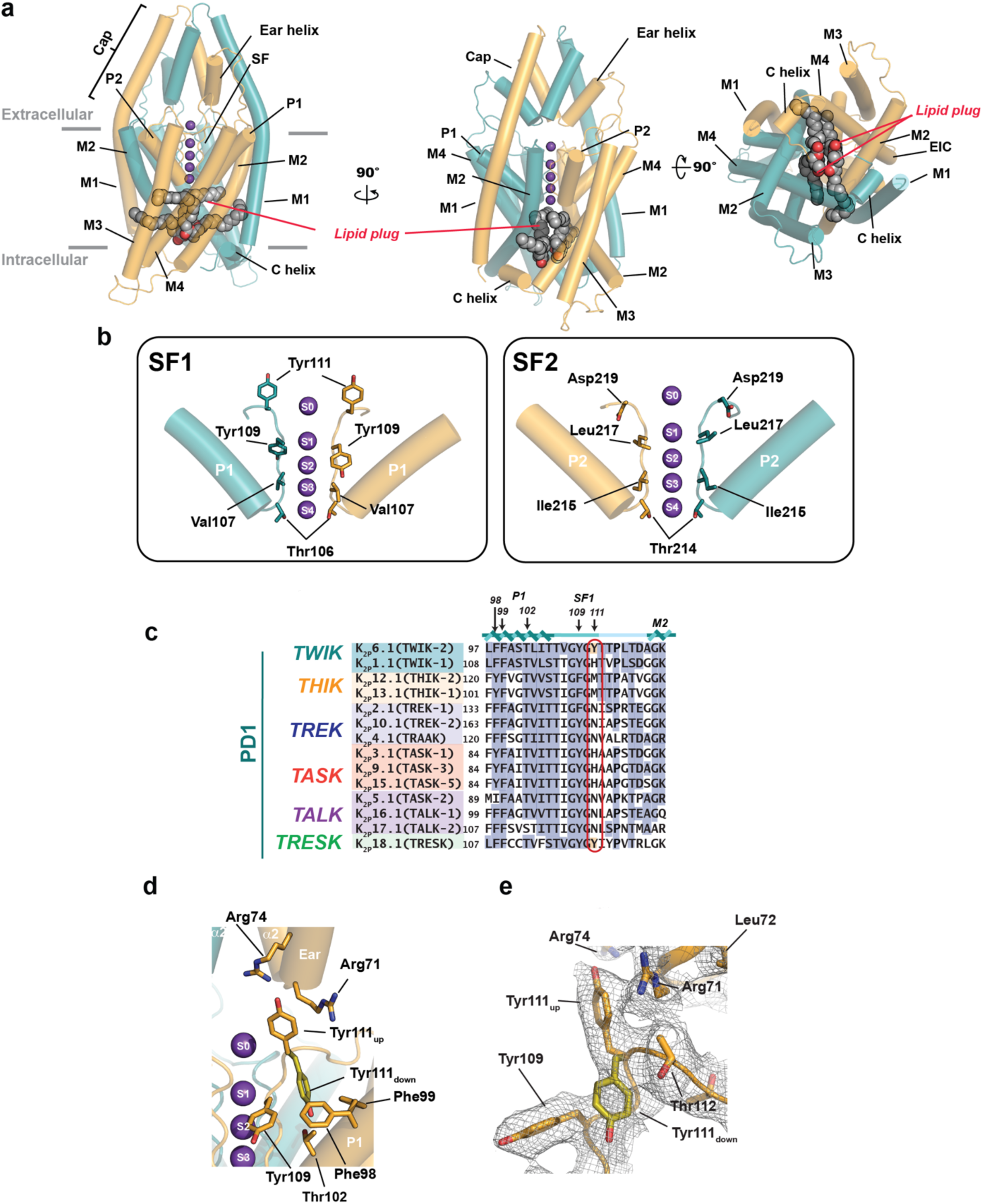
K_2P_6.1(TWIK-2) structural features. **a,** K_2P_6.1(TWIK-2) cartoon diagram (deep teal and bright orange) showing side (left and center) and intracellular (right) views. Lipid plug (grey) is shown in space filling. Grey bars indicate membrane. Potassium ions are shown as purple spheres. **b,** Comparison of K_2P_6.1(TWIK-2) SF1 (left) and SF2 (right) filter conformations. Select matching positions are shown as sticks. **c**, Sequence comparison P1, SF1, and initial segment of M2 from PD1 for K_2P_6.1 (TWIK-2) with other K_2P_s. Red oval indicates Tyr111 site. Labels indicate residues that interact with Tyr111 in the ‘down’ state. Conservation is indicated in blue. Tyr111 and similar residues are shaded red. Sequences are for human: K_2P_6.1(TWIK-2) (GENBANK 4758624); K_2P_ 1.1(TWIK-1) (GENBANK 4504847); K_2P_13.1(THIK-1) (GENBANK 16306555); K_2P_12.1(THIK-2) (GENBANK 11545761); K_2P_2.1(TREK-1) (GENBANK 14589851); K_2P_10.1(TREK-2) (GENBANK 20143944); K_2P_4.1(TRAAK) (GENBANK 15718767); K_2P_3.1(TASK-1) (GENBANK 4504849); K_2P_9.1 (TASK-3) (GENBANK 542133161); K_2P_15.1(TASK-5) (GENBANK 333440483); K_2P_5.1(TASK-2) (GENBANK 333440483); K_2P_16.1(TALK-1) (GENBANK 14149764); K_2P_17.1(TALK-2) (GENBANK 17025230); and K_2P_18.1(TRESK) (GENBANK 32469495). **d**, K_2P_13.1(THIK-1) Try111 environment and interactions. ‘up’ (bright orange) and ‘down’ (olive) Tyr111 conformations are indicated. **e,** Cryo-EM density for the region including Tyr111 (σ=2.5).

**Table 1.**
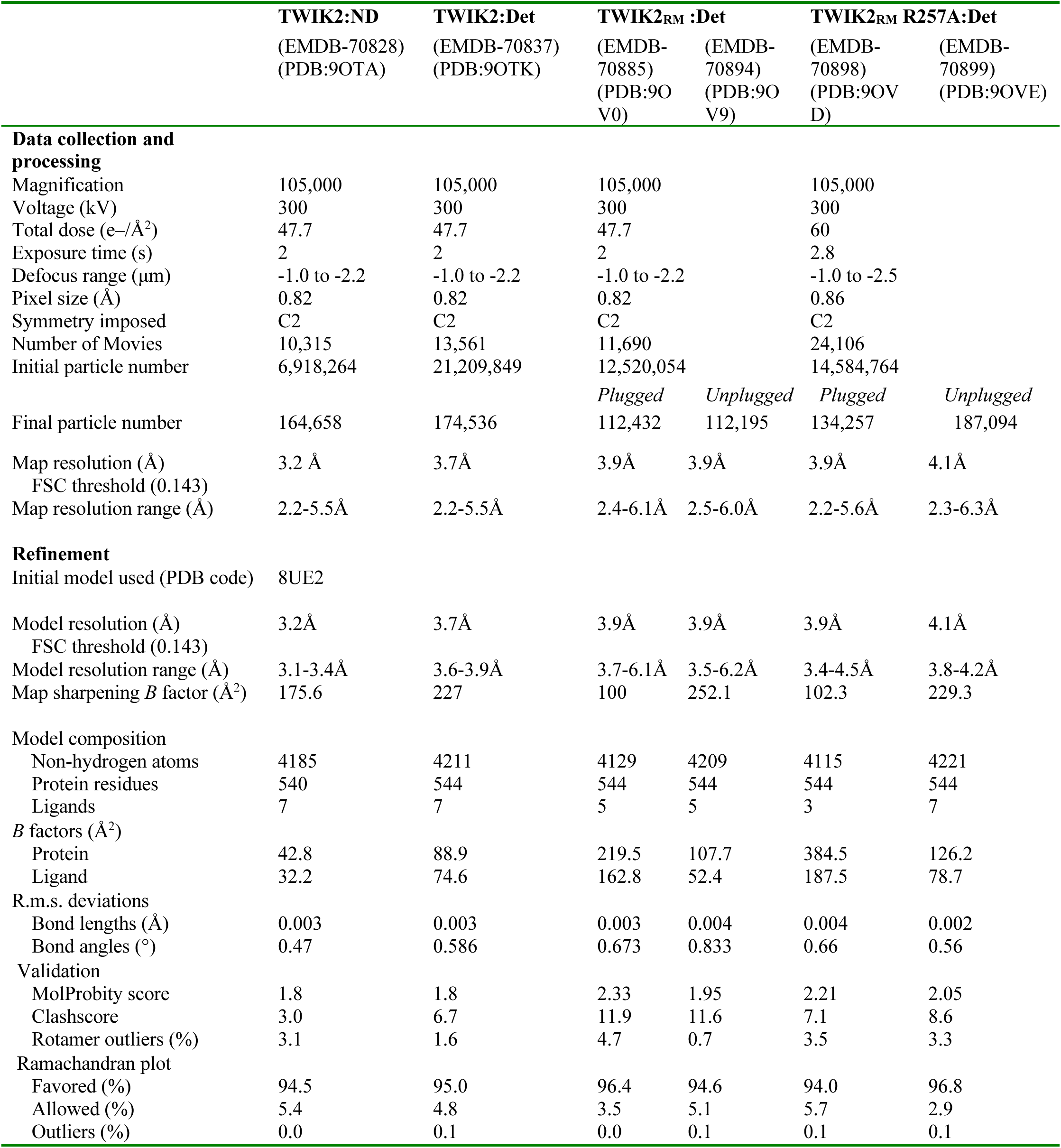
Cryo-EM data collection, refinement and validation statistics.

Consistent with structure solution in high potassium conditions, the TWIK2 selectivity filter shows density for ions at the S1-S4 positions as well as at the extracellular S0 position (Figs. 1b and e, S2b). The selectivity filters of K_2P_ differ from some canonical features seen in other potassium channels. Notably, diverse amino acids occupy the last position of SF1 potassium channel signature selectivity filter sequence ‘TxTTxGYG<u>D</u>’^15,39^ (Fig. 1c), and contrast SF2 where this site is strongly conserved as an aspartic acid (Fig. S7a) as found in homomeric potassium channels^40^. Despite the SF1 sequence variation, in most K_2P_ structures^15^, both SFs adopt essentially the same local conformations in which the sidechain of the residue following the last SF glycine interacts with a network of residues behind the selectivity filter (Fig. S7b), similar to homomeric potassium channels^41^. SF1 in TWIK2 stands out by having a large aromatic residue, Tyr111, following the selectivity filter ‘GYG’ sequence (Fig. 1c). The structure shows that rather than adopting the canonical ‘down’ conformation seen in SF1 from K_2P_1.1 (TWIK1)^4,5^ and K_2P_2.1(TREK-1)^16^ exemplars (Fig. S7b-c) and the equivalent SF2 position (Fig. 1c, Fig. S7b and d), Tyr111 occupies an ‘up’ conformation in which it interacts with Arg71 and Arg74 from the ear helix base (Fig. 1b-d). Inspection of the TWIK2:ND density also showed evidence for a second Tyr111 conformation corresponding to the ‘down’ state (Fig. 1d-e) in which Tyr111 interacts with a pocket formed by P1 helix residues Phe98, Phe99, and Thr102 and SF1 residue Tyr109 (Fig. 1d). The ‘up’ and ‘down’ conformations are essentially Tyr111 rotamer changes that do not require major changes in the surrounding SF structure. Notably, a similar ‘up’ rotamer of the corresponding residue in K_2P_3.1 (TASK-1), His98, was observed in a pH 7.5 structure (Fig. S7e)^42^ contrasting a ‘down’ conformation seen at higher pH^43^. Together, these observations show that the SF1 ‘up’/’down’ rotamer change can occur in different K_2P_ subfamilies. The additional conformational changes observed for the equivalent SF1 position in K_2P_1.1 (TWIK1)^5^ (Fig. S7f) and K_2P_2.1(TREK-1)^25^ under conditions associated with inactivation of the channel C-type SF gate highlight the importance of structural changes this site for K_2P_ function.

The TWIK2 structure also revealed a second notable feature, a pair of branched densities that correspond to the legs two lipids stacked face to face that obstruct the intracellular aqueous cavity (Figs. 1a, 2a, S2a-d). The two legs of each lipid fill the hydrophobic inner cavity above Arg257 and occupy two fenestrations at the interface between P2 and M4 from one subunit and M2 from the other that lead to the membrane bilayer. The upper leg site formed by P2, M2 and M4 is lined by Leu128, Pro132, Met135, Leu246, Met249, Val250, and Leu253. This site corresponds to a site occupied by a single alkyl chain in TWIK1^4^ (Figs. 2b-d, S8a). The lipid lower leg site is formed by M2 and M4 and is lined by Leu136, Leu253, Arg257, Ser260, Thr266, and Leu270 (Figs. 2b-d, S8a). The Arg257 sidechain, a residue conserved in vertebrate TWIK2 (Fig. S5), directly contacts the lipid glycerol backbone (Figs. 2c, S8a). Except for Arg257, the lipid plug contacting residues are conserved between TWIK1 and TWIK2 (Fig. S8b-c).

**Figure 2.**
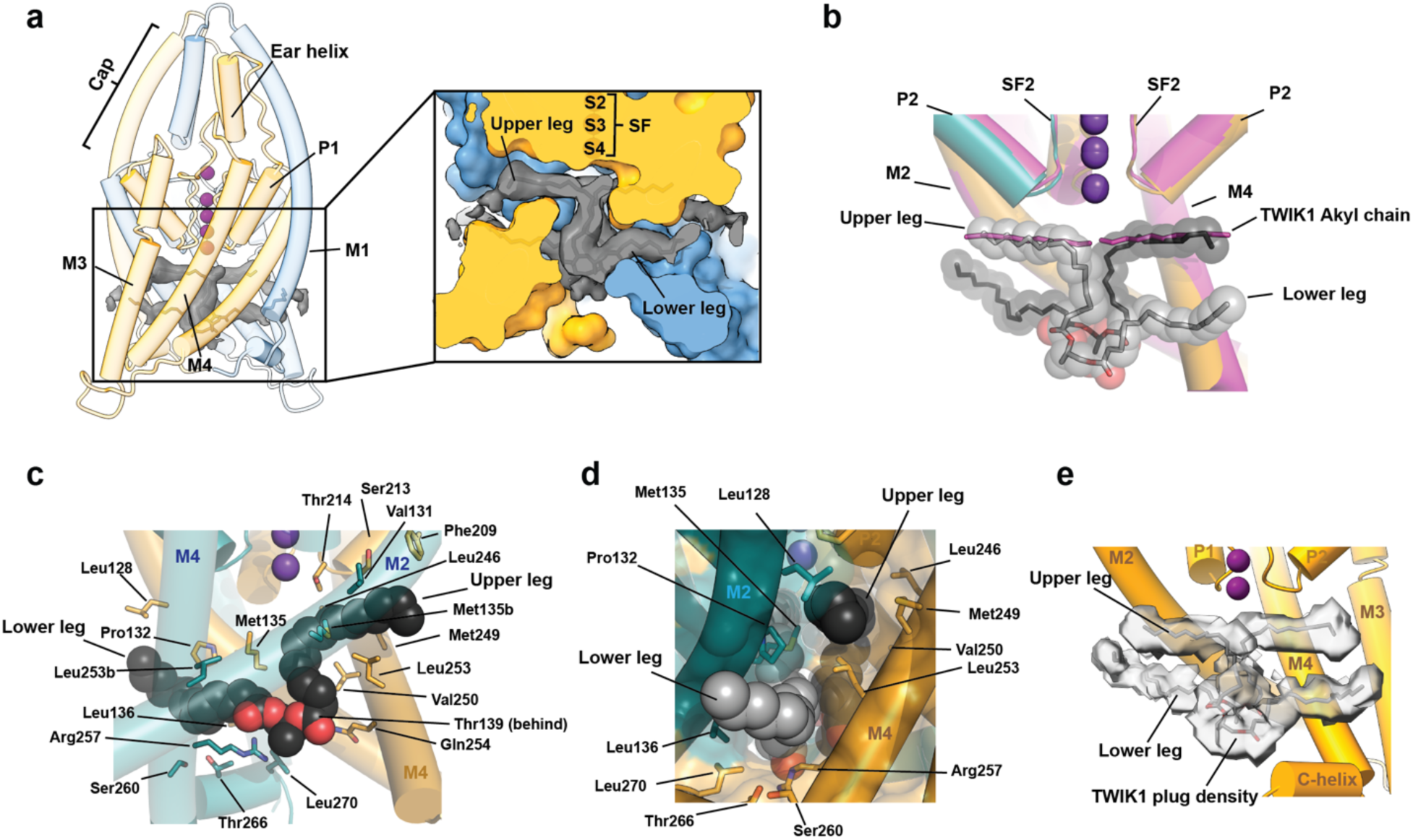
K_2P_6.1(TWIK-2) lipid plug blocks the central cavity. **a,** K_2P_6.1(TWIK-2) cartoon diagram (deep teal and bright orange). Cryo-EM density for lipid plug (grey) is shown (σ =3.0). Inset shows slice through space-filling model of the channel, lipid plug cryo-EM density and stick representation of lipid plug. **b,** Superposition of K_2P_6.1(TWIK-2) (deep teal and bright orange) and K_2P_1.1(TWIK-1) (magenta) (PDB:3UKM)^4^. K_2P_6.1(TWIK-2) lipid plug (grey and black) is shown as sticks and semi-transparent space filling. Upper and lower legs for one lipid are indicated. TWIK1 Alkyl chain^4^ is shown as sticks. **c,** K_2P_6.1(TWIK-2) lipid plug interactions. One lipid plug chain is shown (black, space filling). Contacting residues from the two K_2P_6.1(TWIK-2) subunits (deep teal and bright orange) are shown as sticks. **d,** View from the center of the bilayer towards the upper and lower leg binding sites. Lipid plug chains are grey and black and shown in space filling. Interacting residues are shown as sticks. **e,** Lipid plug density (1.9σ) from K_2P_1.1 (TWIK-1) at pH5.5 (EMDB 25169)^5^. Model shows a single K_2P_6.1(TWIK-2) subunit and lipid plug (grey).

The presence of the lipids in the channel cavity prompted us to ask whether these cavity-filling molecules originated from the nanodisc reconstitution step or were present from an earlier step in channel purification. Hence, we determined the structure of TWIK2 in detergent at 3.7Å (local resolution to 2.2Å) (TWIK2:Det) (Figs. S9-S10, Table 1). TWIK2:Det was essentially identical to TWIK2:ND (RMSD_Cα_=0.44Å) having the same features, including an ‘up’ Tyr111 (Fig. S10a) and ions at sites S0-S4 of the selectivity filter. Importantly, the structure shows density corresponding to the two lipid molecules forming the lipid plug (Fig. S10a), demonstrating that these entities do not originate from the lipids used for the nanodisc reconstitution but come from the cells used to produce the protein sample.

The TWIK1 structure^4^ also has a fenestration below the upper leg site^13^. Comparison with the lipid plug from our TWIK2 structure shows that this cavity matches the position of the plug lower leg (Fig. S8d), suggesting that TWIK1 can accommodate lipids using theTWIK2 bi-legged lipid plug binding pose. Prompted by this structural similarly, we inspected previously reported cryo-EM densities for TWIK1^5^ for evidence of the lipid plug. Remarkably, the density map of TWIK1 at pH 5.5, a structure thought to represent an inactivated channel, shows a bifurcated density in the channel central cavity that is well matched by the TWIK2 lipid plug (Fig. 2e). The map for the TWIK1 pH 7.4 structure, in which the upper site was proposed to be empty of the alkyl chain shows a weaker version of a similar density (Fig. S8e). Thus, the lipid plug binding mode appears to be shared among members of the TWIK subfamily. These observations suggest a revised interpretation of the origin of the alkyl chain density in the upper site of TWIK1 and indicate that it is from one of two legs of a lipid that blocks the pore as we observe in TWIK2.

The conservation of the residues lining both lipid-occupied cavities, particularly in M4, does not extend to other K_2P_ family members (Fig. S8c). Hence, the architecture that enables this bifurcated type of lipid binding in the channel central cavity appears to be a special feature of the TWIK subfamily. Together, these results highlight the unique binding mode of theTWIK2 plug lipid in which each lipid is anchored in the central cavity by occupying the upper and lower leg sites on opposite sides of the channel.

### A trafficking mutation yields unplugged channels

TWIK2 is notable for poor functional expression unless it is directed to the plasma membrane by mutations in lysosomal targeting motifs^14^. To test whether this localization change might affect channel structure or the lipid plug, we determined the structure of a previously characterized TWIK2 mutant^14,33,34^, denoted TWIK2_RM_ (for ‘Retention Mutant’), that is predominantly found at the plasma membrane as a result of a set of C-terminal tail mutations (I289A/L290A/Y308A)^14^. Initial cryo-EM analysis of TWIK2_RM_ showed one form corresponding to the lipid bound state (Fig. S11). However, based on the functional effects of the TWIK2_RM_ mutations^14^, we decided to pursue further classification using a mask focused on the central cavity to determine if we could identify a form representing a functional channel. This analysis yielded two channel forms at 3.9Å resolution each (Fig. S11, Table 1) that we denoted as TWIK2_RM_-Plugged and TWIK2_RM_-Unplugged due to the presence or absence of the lipid plug. The structure of TWIK2_RM_-Plugged was essentially identical to TWIK2:Det (RMSD_Cα_=0.7Å) (Fig. 3a), showed Tyr111 in the up position (Fig. S12a), and two plug lipids blocking the central channel cavity (Fig. 3b-c, S12a). TWIK2_RM_-Uplugged was also overall very similar to TWIK2:Det (RMSD_Cα_=0.7Å) (Fig. 3d) and had Tyr111 from SF1 in the up position (Fig. S13a). Notably, the plug lipid density was absent (Fig. 3f, S13b). Instead, we found a spherical density just below the S4 position of the selectivity filter that we assigned as the ‘cavity ion’ (Figs. 3d-e, S13a), reminiscent of results seen for a structure of K_2P_4.1 in the absence of its fenestration density^6^. Following the identification of the two forms, we reprocessed the TWIK2:ND and TWIK2:Det data using the same strategy used to obtain plugged and unplugged form of TWIK2_RM_ but failed find a class corresponding to the unplugged form. These data suggest that the strong intracellular retention of the wild-type channel is associated with the lipid plugged form, whereas some fraction of the channel lacks this plug when the channel can reach the plasma membrane. The region bearing the trafficking mutations is not seen in any of our structures, suggesting that this region is disordered. The density for the ions in the selectivity filter of both the TWIK2_RM_ plugged (Fig. S12a) and TWIK2_RM_ unplugged (Fig. S13a) structures suggests differences in ion occupancy at the S0 and S1 sites from the TWIK2 structures. The structural analysis of the trafficking mutant TWIK2_RM_ shows that the channel can exist in two states, one having two lipid plug molecules (TWIK2:ND, TWIK2:Det, and TWIK2_RM_:Det plugged) and one in which access to the central cavity is unhindered (TWIK2_RM_:Det unplugged). Comparison of representative structures for the plugged (TWIK2:ND) and unplugged (TWIK2_RM_:Det unplugged) forms shows that the two plug lipids effectively block ion access to the channel by filling the central intracellular cavity below the selectivity filter (Fig. 3g-h).

**Figure 3.**
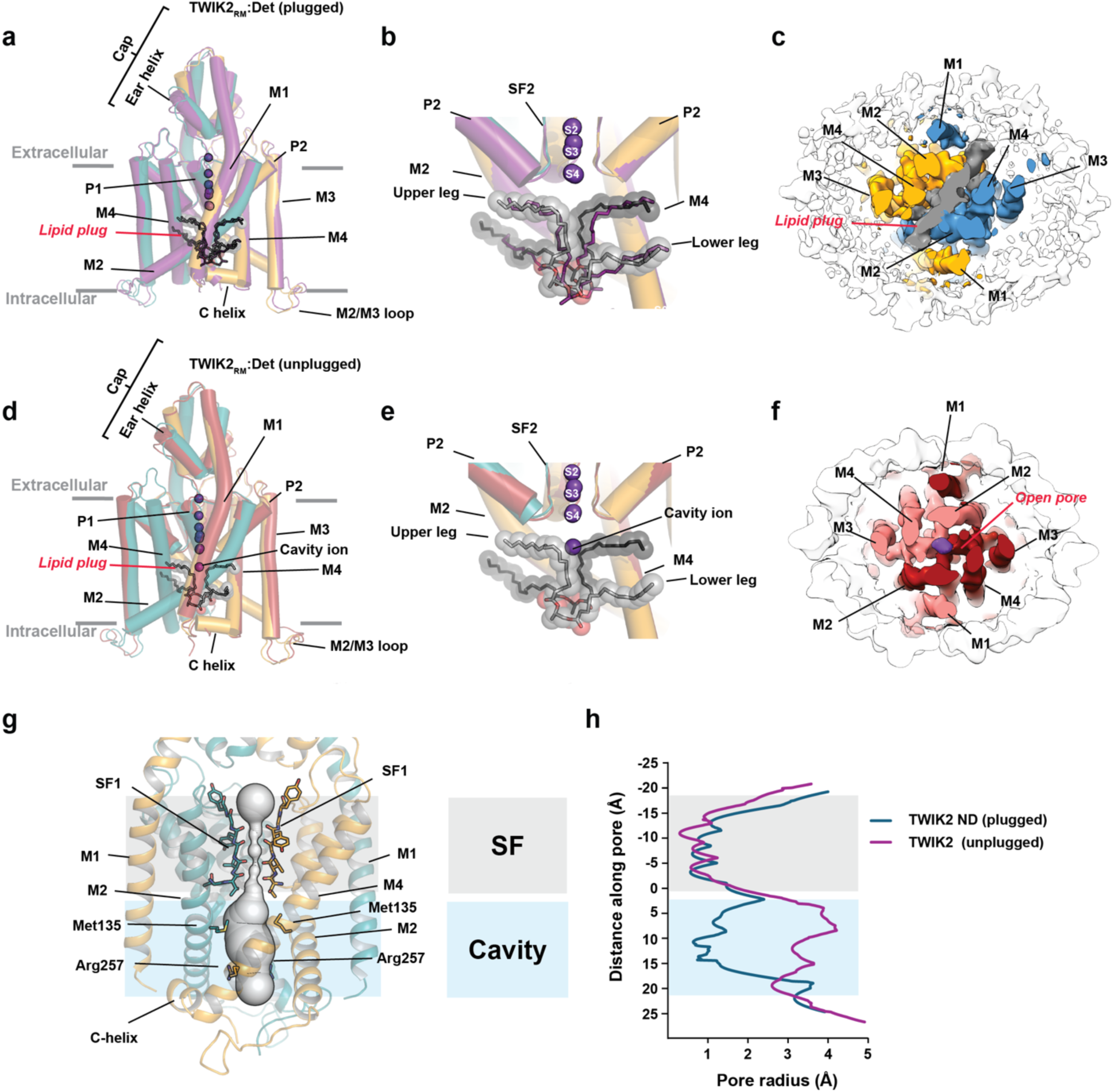
Comparison of plugged and unplugged K_2P_6.1 (TWIK2) structures. **a**, Superposition of TWIK2:ND (deep teal and bright orange) and TWIK2_RM_:Det (plugged) (deep purple) structures. **b,** Close up view of the central cavity from ‘a’. Lipid plugs from TWIK2:ND and TWIK2_RM_:Det (plugged) are shown as spheres (grey) and sticks (deep purple), respectively. **c,** Intracellular view of a slice through the TWIK2_RM_:Det (plugged) density. Subunits are deep teal and bright orange, lipid plug density is grey. **d,** Superposition of TWIK2:ND (deep teal and bright orange) and TWIK2_RM_:Det (unplugged) (firebrick) structures. **d,** Close up view of the central cavity from ‘d’. Lipid plug from TWIK2:ND is shown as spheres (grey). TWIK2_RM_:Det (unplugged) cavity ion is labeled. **f,** Intracellular view of a slice through the TWIK2_RM_:Det (unplugged) density. Subunits are firebrick and light red. **g,** Pore profile of K_2P_6.1(TWIK2) (deep teal and bright orange) calculated using HOLE ^53^. Selectivity filter and key cavity positions are shown as sticks. **h,** K_2P_6.1(TWIK2) pore profiles for plugged (blue) and unplugged (magenta) forms. Selectivity filter (SF) (grey) and cavity (blue) are indicated. in ‘g’ and ‘h’.

### Functional studies highlight the importance of lipid-interacting cavity residues and Tyr111

We set out to test key features identified by the TWIK2 structures. As previously reported^10,14^, TWIK2 did not produce measurable currents in *Xenopus* oocytes using two electrode voltage clamp (TEVC), whereas mutation of the endolysosomal targeting signal (I289A/L290A/Y308A)^14^ (TWIK2_RM_) yielded readily measurable TWIK2 channels (Fig. S14a) that provided a platform for testing the effects of mutants. To test the importance of the lipid plug contacts, we examined the effects of mutations at lipid plug-channel at three places along the channel central axis (Fig. 4a-c), with a focus on mutation to negatively charged residues to destabilize lipid-channel contacts. V131D, located near the base of the selectivity filter, did not make functional channels (Figs. 4b-c, S14a). However, introduction of acidic residues at Met135 (M135D) and Arg257 (R257E) resulted increased basal activity with respect to TWIK2_RM_, as did replacement of Arg257 by alanine (R257A) (Fig. 4b-c, S14a). Comparison of ion selectivity by following the reversal potential as a function of external potassium showed no significant differences in the functional mutants from TWIK2_RM_ (Fig. S14b). The increased activity observed for mutation at sites that interact with the blocking lipids is consistent with the idea that removal of the lipid plug is key to channel function. Further the fact that introduction of a negative charge and truncation to alanine at the Arg257 site increases activity points to an important role for this position in TWIK2 function.

**Figure 4.**
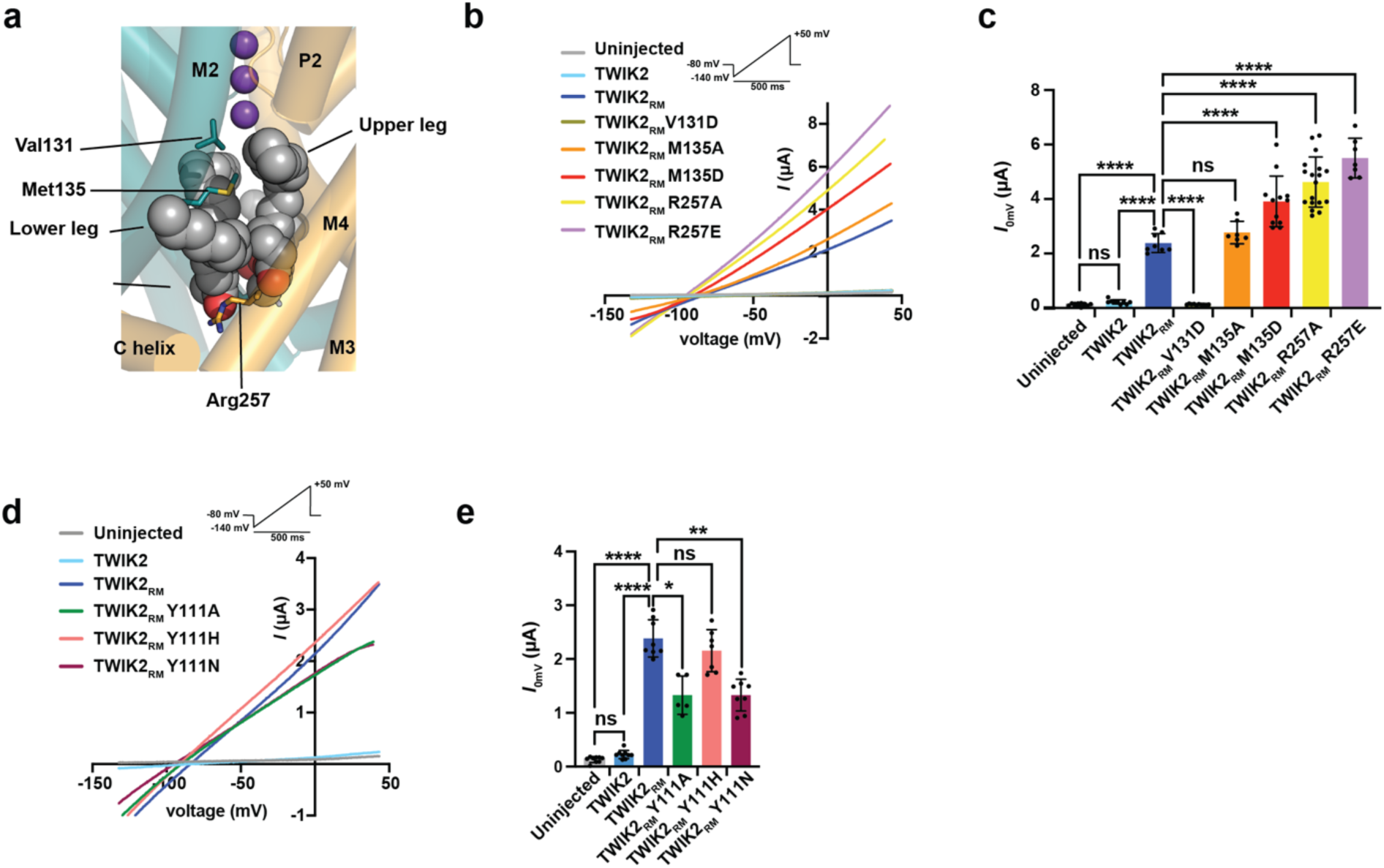
K_2P_6.1(TWIK2) cavity and filter sites affect function. **a,** View of the K_2P_6.1(TWIK2) lipid plug from center of the membrane. Lipid plug is shown as grey spheres. Key interacting residues are shown as sticks. Potassium ions are purple spheres. **b,** Exemplar current-voltage responses from TEVC *Xenopus* oocytes expressing the indicated channels and cavity mutants. Uninjected (grey), TWIK2 (sky blue), TWIK2_RM_ (blue), TWIK2_RM_ V131D (olive), TWIK2_RM_ M135A (orange), TWIK2_RM_ M135A (red), TWIK2_RM_ R257A (yellow), and TWIK2_RM_ R257E (lavender). Inset shows protocol. **c,** Effects of mutants on K_2P_6.1(TWIK2) basal activity at 0 mV. **** p <0.0001, *** p <0.001, ** p < 0.01, * p = 0.01 -0.05. Statistical analysis was performed using a One-Way ANOVA and multiple comparisons were performed with Tukey’s test. Error bars are S.E.M.. **d,** Exemplar current-voltage responses from TEVC *Xenopus* oocytes expressing the indicated channels and mutants. Uninjected (grey), TWIK2 (sky blue), TWIK2_RM_ (blue), TWIK2_RM_ Y111A (green), TWIK2_RM_ Y111H (salmon), and TWIK2_RM_ Y111N (maroon). Inset shows protocol. **e,** Effects of mutants on K_2P_6.1(TWIK2) basal activity at 0 mV. **** p <0.0001, *** p <0.001, ** p < 0.01, * p = 0.01 -0.05. Statistical analysis was performed using One-Way ANOVA and multiple comparisons were performed with Tukey’s test. Error bars are S.E.M..

The fact that Tyr111 is the largest residue seen at this filter position (Fig. 1c) and was observed in both up and down conformations prompted us to test effect of replacing this residue with smaller, more commonly observed amino acids or with alanine. Replacement of this position with the most common residue found at this site in other K_2P_s, asparagine (Y111N) (Fig. 1c) caused a reduction in basal current, as did replacement with alanine (Y111A). By contrast, Y111H yielded basal currents similar to wild type. Despite the fact that the mutation site is in the selectivity filter, none of these changes affected responses to external potassium (Fig. S14d) indicating that the selectivity of the channels remained intact. Notably, all three changes resulted in channels that showed inactivation (Fig. S14 a and c) supporting the importance of Tyr111 in the function of the K_2P_6.1 (TWIK2) selectivity filter gate.

### R257A mutation changes lipid plug position

To investigate the structural consequences of perturbing lipid-coordinating residues (Fig. 4c), we determined the structure of TWIK2_RM_ R257A, a mutant that increases TWIK2_RM_ basal currents (Fig. S15). As with TWIK2_RM_, cryo-EM analysis of single particles of TWIK2_RM_ R257A using focused classification on the central cavity yielded two channel classes, one having the lipid plug, TWIK2_RM_ R257A-plugged (3.9Å resolution), and one in which the plug was absent, TWIK2_RM_ R257A-unplugged (4.1Å resolution) (Figs. S15-S17, Table 1). The overall structures of both TWIK2_RM_ R257A forms were similar to the TWIK2:ND structure (Fig. 5a-d) (RMSD_Cα_=0.82Å). Both had Tyr111 in the up position (Figs. S16a and S17a). TWIK2_RM_ R257A-plugged showed clear ion density at selectivity filter sites S0-S4 (Fig. S16a), whereas there were fewer well-defined ions in the TWIK2_RM_ R257A-unplugged form (Fig. S17a). Strikingly, in the TWIK2_RM_ R257A-plugged form, the lipid plug resides ∼5Å closer to the intracellular opening of the channel than in the other plugged TWIK2 structures (Figs. 5a-b, and S18a). Because of this lower position, ∼four upper site methylene groups and ∼ten lower site methylene groups are pulled into the central cavity leaving a smaller portion of the upper and lower legs in their respective binding sites (Fig. 5a-b, and S18b). This lower pose also reveals density contacting the C-helices that is consistent with phosphatidylethanolamine head groups (Fig. 5a-b, S18a). The observation that the R257A affects lipid plug position indicates that this residue has a key role in lipid plug binding and suggests that the functional effects from mutations at this site (Fig. 4c) are rooted in alteration of lipid plug binding. Importantly, the fact that we can observe positional changes in the lipid plug provides corroborating evidence regarding its assignment and role as a crucial component of the TWIK2 structure.

**Figure 5.**
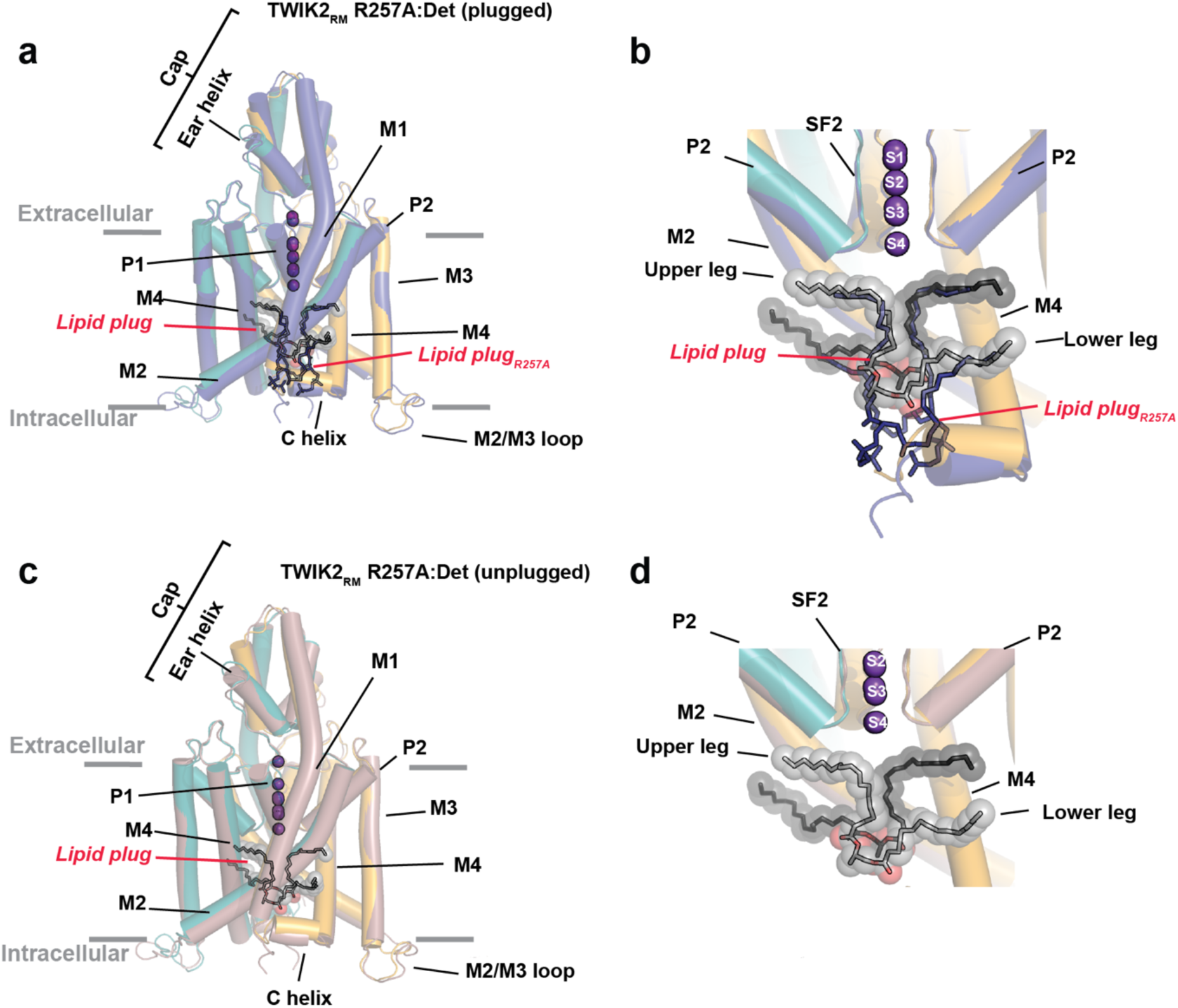
Comparison of plugged and unplugged K_2P_6.1 (TWIK2) R257A structures. **a,** Superposition of TWIK2:ND (deep teal and bright orange) and TWIK2_RM_ R257A:Det (plugged) (deep blue) structures. **b,** Close up view of the central cavity from ‘a’. Lipid plugs from TWIK2:ND and TWIK2_RM_ R257A:Det are shown as spheres (grey) and sticks (deep blue), respectively. **d,** Superposition of TWIK2:ND (deep teal and bright orange) and TWIK2_RM_ R257A:Det (unplugged) (dirtyviolet) structures. **d,** Close up view of the central cavity from ‘c’. Lipid plug from TWIK2:ND is shown as spheres (grey).

## Discussion

A puzzling aspect of many ion channel structures has been the identification of fenestrations from the central cavity that contain lipid or lipid-like densities that could represent elements that interfere with the passage of ions through the channel central cavity^4,6,7,44^. In particular, this phenomenon has been observed for a number of K_2P_s, including K_2P_1.1 (TWIK1)^4,5^, K_2P_4.1 TRAAK^6,7^, K_2P_2.1 (TREK1)^12^, and K_2P_10.1 TREK2^8^. However, in each case the reported densities only define a single alkyl chain^4,6^ leaving open the possibility that its origin could be detergents required for channel purification rather than a lipid^4,12^. One interpretation of this density is that it represents a mechanism in which horizontal lipid access from the bilayer acts as a gating mechanism^5,6^; however, this issue remains controversial^13,27,45^, particularly as simulations indicate that even though lipid tails can enter the cavity from the bilayer they do not block the pore^13,27^.

Here, we provide evidence for TWIK2 block by a pair of two-legged lipids, the lipid plug, in which each lipid leg fills a separate binding site that extends towards the bilayer and the lipid headgroup points towards the inner cytoplasmic face in an orientation that matches that of bilayer inner leaflet lipids. The upper leg site overlaps with the position assigned as single alkyl chain in the initial structure of K_2P_1.1 (TWIK1)^4,5^ framed by P2, M2, and M4 (Fig. 2b). The lower leg passes through a gap on the opposite side of the channel between M2, M4, and the C-helix. In this binding mode, each of the plug lipids spans the inner cavity. Given the high similarity between TWIK1 and TWIK2 structures (Fig. S3a-c), excellent overlap of the upper leg site (Fig. 3b), and presence of the lower cavity in TWIK1^13^ (Fig. S8d), our data suggest that the lipid block we observe occurs in both channel types. This interpretation is strengthened by the fact that cryo-EM maps from TWIK1 in an inactive (pH 5.5) and active (pH 7.4) forms have a bifurcated density in the central cavity that is well matched by the lipid plug structure derived from our TWIK2 studies (Figs. 2e and S8e). Thus, previous observations of density in the upper leg site may represent incomplete density for similar lipid plug structures arising from lipid dynamics. Together, these data reveal a lipid binding mode that has not been noted previously and offer a reinterpretation of previous findings regarding how lipids bind TWIK channel fenestrations. Structural analysis of the R257A mutant that removes interactions with the blocking lipids leads to a repositioning of the lipid plug and suggests an important role for the conserved Arg257 residue in TWIK2 channels. It is worth noting that TWIK1 and TWIK2 have narrower inner pores than channels from the TREK subfamily, suggesting that the lipid plug structure is a distinct feature of the TWIK subfamily.

Ion concentrations inside various intracellular organelles vary and there are a growing number of ion channels whose activity affects diverse intracellular compartments^46^, including TWIK2^47^. Because every ion channel has the potential to perturb ion flux, it is essential that these protein pores open only in the right place at the right time. The majority of channels in the VGIC superfamily that are activated by voltage or ligands have an intracellular gate that blocks ion passage, rendering them inactive in the absence of a stimulus^48^. This structural barrier avoids openings in the wrong subcellular compartment. Although their activity can be tuned by various types of inputs^2,15^, the fundamental property of K_2P_s is that they are leak channels, lacking the strongly closed state of their VGIC superfamily relatives^2,15^. Consequently, their basal activity gives K_2P_s the capacity to influence ion flux across both internal and external membranes as they transit through various cellular compartments during their life cycle. Such basal activity also has the potential to inadvertently lead to aberrant functional effects if it is unregulated. We propose that the propensity of TWIK channels to be plugged by lipids represents a regulatory mechanism that contributes to the ability of a cell to control K_2P_ function (Fig. 6). Such changes in activity may be linked processes in which channel activation involves redistribution of TWIK2 channels from intracellular organelles to the plasma membranes as has been suggested during macrophage inflammasome activation^30,32–34^. Together, our observations suggest the hypothesis that TWIK family activation is a ‘right place, right time’ process in which the channels remain plugged until they arrive at the cellular destination where their activity is required.

**Figure 6.**
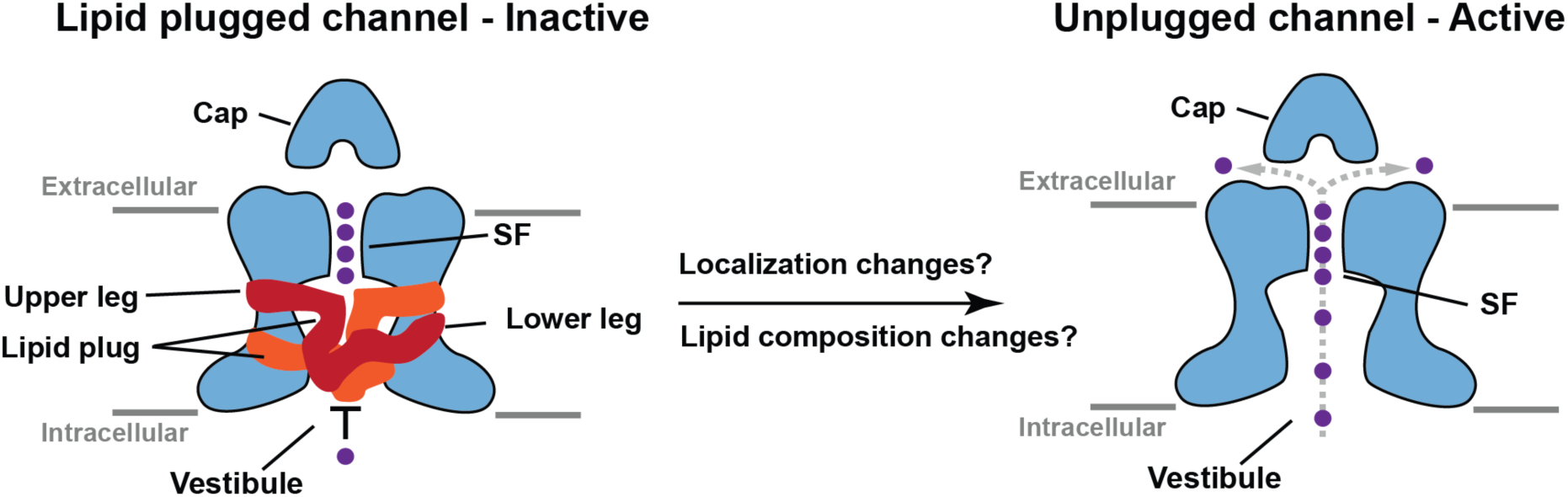
Model of K_2P_6.1(TWIK2) regulation. Lipid plug prevents ion passage. Removal of plug in response to cellular localization changes, changes in the membrane lipid composition, or other factors yields channels that can conduct ions.

Although our data establish that the channel can exist in both plugged and unplugged forms, how the lipid plug moves in and out is unclear. The presence of the plug is incompatible with ion movement through the central cavity (Fig. 3h). Hence, its removal from the cavity is essential to yield a functional channel. Changes in the positions of the M4 and the C-helix would offer a potential exit path as such motions would alter the size of the lateral gap facing the bilayer that forms the most natural way for lipid egress. Whether such conformational transformations constitute an autonomous mechanism driven by changes in lipid bilayer properties or requires the intervention of factors that facilitate lipid removal remains an open question in need of investigation. Wetting-dewetting of the narrow hydrophobic inner pore has been proposed as an important regulatory mechanism for TWIK channels that can be influenced by occupation of the lateral fenestration by lipid tails^13,49,50^. As such, there may also be some role for the competition between water and the lipid plug. Further, a recent structure of TWIK2 that was reported while this work was in preparation suggests that the density previously assigned to a single alkyl chain that we assign to the upper leg of the plug can be displaced by pimozide, a drug that modulates TWIK2, raising the possibility of competition between the lipid plug and small molecule modulators of K_2P_ function^51^.

Besides the new insights into mechanisms of lipid modulation, the TWIK2 structure highlights the unusual divergence from the canonical selectivity filter sequences afforded by the heteromeric nature of the K_2P_ SF^15,39^. The presence of a large, non-canonical residue at Y111, a site usually occupied by smaller residues (Fig. 1c) shows two conformations. The ‘up’ conformation is predominant in our structures but is also accompanied by a ‘down’ conformation in the lipid nanodisc complex that matches the pose found for the equivalent position in other K_2P_s (Fig. S7b-d). These ‘up’/’down’ conformations match those of a regulatory histidine, His98, in K_2P_3.1 (TASK-1)^42,43^ (Fig. S7e), and together with the functional effects of Tyr111 mutations (Fig. 4d-e) highlight the role of this site in K_2P_ function and the importance of the K_2P_ C-type selectivity filter gate^22,23,26,52^ for TWIK2 function. The observation that the ‘up’/’down’ rotamer change can occur in different K_2P_ subfamilies together with additional conformational changes reported for the equivalent SF1 position in K_2P_1.1 (TWIK1)^5^ (Fig. S7f) and K_2P_2.1(TREK-1)^25^ under conditions associated with inactivation of the channel C-type SF gate highlight the functional importance of structural changes at this site.

Structures ofTWIK2 in lipid and detergent environments shows that this channel conforms to the overall architecture found throughout the K_2P_ family and is most similar to the other structurally characterized member of the TWIK subfamily, TWIK1. Nevertheless, the TWIK2 structure revealed three new elements: an extra helix in the extracellular cap domain, the ear helix, that connects the cap with the loop leading to the first pore helix (P1), conformational variability at a position in SF1 that is linked to C-type gating in other K_2P_s^5,25,42,43^, and the lipid plug. Understanding the roles of these elements in K_2P_6.1 (TWIK2) function will be important for dissecting the biological functions of this channel including how it affects macrophage NLRP3 inflammasome activation^30,32–34^. Our reinterpretation of how lipids affect TWIK channels by acting as a plug that binds to two lateral gaps in the channel pore sets a new framework for addressing how ion channel-lipid interactions control function that may have general relevance for other ion channel classes.

## Acknowledgements

We thank A. Cassago and P. Pascual at the S2C2 Cryo-EM Center and D. Bulkley and G. Gilbert at the UCSF Cryo-EM facility for help with microscope handling and data acquisition, M. Grabe for helpful discussions, and K. Brejc and M. Grabe for comments on the manuscript. This work was supported by NIH grant R01-MH093603 to D.L.M.

## Author Contributions

A.M., S.N., and D.L.M. conceived the study and designed the experiments. A.M. expressed and purified the proteins, prepared the cryo-EM samples, collected and analyzed the cryo-EM data, built and refined the models. S.N. collected and analyzed the two-electrode voltage clamp data. A.M., S.N., and D.L.M. analyzed the data. D.L.M. provided guidance and support. A.M., S.N., and D.L.M. wrote the paper.

## Competing interests

The authors declare no competing interests.

## Materials and Methods

### References

### Materials and Methods

#### Expression and purification of K2P6.1WT and mutants

The gene for a construct of the human K_2P_6.1 (TWIK-2) (Uniprot ID: Q9Y257), K_2P_6.1_WT_, spanning residues 1-313 followed by a 3C protease cleavage site, monomeric enhanced green fluorescent protein (mEGFP), and a His_8_ tag was cloned into a modified pFastBac expression vector in which the polyhedrin promotor was replaced by a mammalian cell active CMV promotor^54^. Expression vectors for TWIK-2_ERM_ (I289A, L290A and Y308A) and TWIK-2_ERM_ R257A mutants were made on this background. These vectors were used for recombinant bacmid DNA generation using chemically competent DH10EmBacY (Geneva Biotech) cells. These bacmids were then used to transfect *Spodopetera frugipdera* (SF9) cells to make baculoviruses for the constructs of interest^55^.

Expi293F cells (Gibco A14528) were grown to 2.5–3 x 10^6^ cells mL^-1^ at 37 °C supplemented with 8 % CO_2_ shaking at 120 RPM before transduction with 10 % (v/v) baculovirus stock of K_2P_6.1_WT_ or mutants. 12 – 14 h after transduction, 10 mM sodium butyrate was added to the culture to enhance protein expression^56^. Flasks were then moved to 30 °C for 48h before harvesting the cells using centrifugation at 2300*g* for 20 min. The pellet was gently washed with Dulbecco’s phosphate buffered saline (Gibco 14190144) to remove leftover culture media, centrifuged (3500 x g for 20 min) to recover the cell pellet, and then flash frozen with liquid nitrogen for storage at – 80°C.

The pellet from 1.8 L culture was resuspended in 50 mL hypotonic buffer (20 mM KCl, 10 mM Tris-HCl pH 8.0, phenylmethylsulfonyl fluoride (PMSF), 0.1 mg mL^-1^ DNase1, supplemented with Pierce protease inhibitor (Thermo Fischer Scientific) and was gently stirred on a Mono Direct Variomag magnetic stirrer (Thermo Fischer Scientific) at 4 °C for 30 min. The membranes were isolated by ultracentrifugation at 185,500*g* for 30 mins. Using a 16 G Precision Glide needle and 30 mL syringe, the membrane pellet was homogenized in 200 mL solubilization buffer (buffer S) (100 mM KCl, 50 mM Tris-HCl pH 8.0, supplemented with 1 mM PMSF, 0.1 mg mL^-1^ DNase1, 1 % (w/v) n-Dodecyl-β-D-Maltopyranoside (β-DDM), and Pierce protease inhibitor. Membrane solubilization and protein extraction was performed by gently stirring at 4°C for 3 h followed by ultracentrifugation at 185,500*g* for 40 min to remove cell debris. 5ml of anti-GFP nanobody conjugated resin conjugated CNBr Sepharose beads (GE Healthcare, #17-0430-02)^57^ pre-equilibrated with 200 mM KCl, 0.01% β-DDM, 20 mM Tris-HCl pH 8.0 buffer (buffer C) was added to the supernatant after the ultracentrifugation and incubated at 4°C with gentle rotation on a Boekel 260200 Orbiton platform rotator. Subsequently, the resin was collected on a gravity flow column and washed with 10 column volumes (CV) of buffer A (200 mM KCl, 20 mM Tris-HCl pH 8.0, 0.1% β-DDM), followed by another wash step with 10 CV of buffer B (200 mM KCl, 20 mM Tris-HCl pH 8.0, 0.05% β-DDM), and then 10 CV of buffer C (200 mM KCl, 20 mM Tris-HCl pH 8.0, 0.01% β-DDM). On column cleavage of the affinity tag was achieved by incubating the resin overnight at 4 °C without any shaking with buffer C supplemented to contain 200 mM KCl, 1 mM EDTA, and 3C protease at ratio of 50:1 resin volume:protease volume. The cleaved sample was collected in 50 mL falcon tube, and the resin was subsequently washed with 2 column volumes (CV) of buffer C. The wash was also collected in the same 50 mL falcon tube. This purified sample was concentrated using Amicon Ultra-15 centrifugal unit with 100 kDa cutoff and applied to a Superdex 200 increase column pre-equilibrated with buffer C. Peak fractions were analyzed by SDS-PAGE and concentrated to 2 mg mL^-1^ for cryo-EM sample preparation using an Amicon Ultra-0.5 ml centrifugal unit with 100 kDa cutoff.

For nanodisc (ND) reconstitution, Brain Extract Polar lipid (Avanti polar lipids #141101P) was prepared by dissolving in 100 mg of lipid in 0.2 mL chloroform, drying under nitrogen gas, and followed by lyophilization. The lyophilized lipid was dissolved by sonication in 1mL of lipid buffer (200 mM KCl, 50 mM Tris pH 8.0, 0.5 % β-DDM). TWIK2 from the size-exclusion chromatography step, purified MSP1E1^38^, and prepared lipid stock were incubated together at 1:5:250 molar ratio (nanodisc reconstitution mix) for 1 h in a 1.5mL Eppendorf microcentrifuge tube at 4 °C. Bio-beads SM-2 adsorbents (Bio-rad #1528920) were prepared by washing 200 mg bio-beads with 1mL methanol, followed by 1mL water and then with 1mL buffer N (200 mM KCl, 20 mM HEPES pH 7.4). 100 mg of bio-beads equilibrated with buffer N were added to 1 mL nanodisc reconstitution mix and incubated for 1 h gently rotating at 4 °C. After 1hr the reconstitution mix was transferred to a clean 1.5 mL Eppendorf tube and incubated with another 100 mg of equilibrated bio-beads and was rotated gently at 4 °C overnight. The supernatant from the reaction mixture was injected onto Superose 6 increase 10/300 column pre-equilibrated with buffer N. Peaks were analyzed by SDS-PAGE and concentrated to 3 mg mL^-1^ for cryo-EM sample preparation using an Amicon ultra-0.5 mL centrifugal unit with 100 kDa cutoff.

Twik-2_RM_ and Twik-2_RM_R257A were expressed and purified by following identical methods as used for Twik-2_WT_. Following the final Superose 6 Increase 10/300 GL purification step, purified samples were concentrated using Amicon ultra-0.5 mL centrifugal units to 3 mg mL^-1^ for cryo-EM sample preparation.

#### Cryo-EM sample preparation and Data collection

3.5 µL of concentrated protein sample was applied to a glow-discharged (30 s at 15 mA) grid (Quantifoil Au R1.2/1.3 300 mesh) and after a wait time of 5 s, grids were blotted once for 2 – 6 s (4 °C and 100 % humidity) using a FEI Vitrobot Mark IV (Thermo Fisher Scientific) and plunge-frozen in liquid ethane. The grids were screened using Serial EM ^58^ on 200 kV Talos Arctica cryo-TEM (Thermo Fisher Scientific) equipped with K3 direct detector camera (Gatan) at the University of California, San Francisco (UCSF) EM facility. Movies were collected using Serial EM ^58^ and EPU (Thermo Fisher Scientific, https://www.thermofisher.com/us/en/home/electron-microscopy/products/software-em-3d-vis/epu-software.html) data acquisition software on 300 keV FEI Titan Krios (Thermo Fisher Scientific) at UCSF and SLAC National Accelerator Laboratory, respectively. Both microscopes were equipped with a K3 direct-electron detector and post-BioQuantum GIF energy filter (Gatan), set to a slit width of 20 eV. Super-resolution counting mode was used at a nominal magnification of x105,000 with a pixel size of 0.43 Å (SLAC) and 0.42 Å (UCSF). A nominal defocus range between -1.1 to -2.5 µm was used with total dose of 60 e^-^ / Å^2^, having total exposure time of 2.8 s at SLAC and -1.1 to -2.2 µm defocus range was used with total dose of 47.7 e^-^ / Å^2^, having total exposure time 2 s at UCSF EM facility.

#### Structure determination (image processing, model building and refinement)

A total of 10,315, 13,561, 11,690 and 24,106 movies were collected for TWIK2:ND, TWIK2:Det, TWIK2_RM_:Det and TWIK2_RM_R257A:Det samples, respectively. Cryo-EM data processing was performed using cryoSPARC (version 4.5)^59^ and Relion (v5)^60^. Patch motion correction (2x binned) and patch CTF estimation were performed before manually curating the exposures by ice-thickness, CTF-fit resolution. Selected micrographs were used for reference-free circular blob picker (diameter 60-120 Å) for particle picking followed by extraction at a box size of 260 Å. Several rounds of 2D classification were performed and further cleaning was done by categorizing particles by 3D *Ab initio* reconstruction. Finally, non-uniform and local refinement were performed to improve resolution and map quality. Datasets corresponding to Twik-2_RM_ and Twik-2_RM_R257A yielded plugged conformations after non-uniform refinement. The refined particle sets for each of these datasets were subjected to 3D classification using a focused mask corresponding to the lipid-plug density obtained in non-uniform refinement. The 3D classification segregated the total particle stacks in two major groups (plugged and un-plugged). These two groups were further subjected to local refinement to obtain the final maps corresponding to Twik-2_RM_ (plugged and un-plugged) and Twik-2_RM_R257A (plugged and un-plugged). A similar approach, generating lipid plug mask using the density observed in the vestibule region and using it for focused 3D classification, was attempted to find un-plugged conformations in the TWIK2:ND and TWIK2:Det datasets. However, this approach did not generate any 3D class without a lipid plug for these two datasets, indicating an absence of the an unplugged class.

A preliminary model based on K_2P_2.1 (TREK-1)(PDB ID: 8UE2) ^61^ was converted to poly-alanine and used to dock as a backbone in the density using phenix.dock_in_map (v1.21.2) ^62^. Multiple rounds of iterative model building and refinement was performed using Coot (v0.9.8.95)^63^ and phenix.real_space_refine^64^, respectively. Final geometry validation statistics were calculated by MolProbity^65^. Figures were prepared and model comparisons were performed using UCSF ChimeraX (version 1.15) ^66^ and the Pymol package (http://www.pymol.org/pymol). Contact analysis was performed using LIGPLOT^67,68^.

#### Two-electrode voltage-clamp (TEVC) electrophysiology

*Xenopus laevis* oocytes were harvested according to UCSF IACUC Protocol AN193390 and digested using collagenase (Worthington Biochemical Corporation, #LS004183, 0.8-1.0 mg mL^-1^) in Ca^2+^-free ND96 (96 mM NaCl, 2 mM KCl, 3.8 mM MgCl_2_, 5 mM HEPES pH 7.4). Prior to RNA injection, oocytes were maintained at 4 °C in ND96 (96 mM NaCl, 2 mM KCl, 1.8 mM CaCl_2_, 2 mM MgCl_2_, 5 mM HEPES pH 7.4) supplemented with antibiotics (100 units ml^-1^ penicillin, 100 µg ml^-1^ streptomycin, 50 µg ml^-1^ gentamicin) and used for experiments within one week of harvest.

Human K_2P_6.1 (TWIK-2) (Uniprot ID: Q9Y257) was subcloned into a previously reported pGEMHE/pMO vector^22^. Site directed mutations were made using Inverse PCR ^69^ with fully overlapping primers. mRNA for oocyte injections was prepared from linearized plasmid DNA (linearized with AflII) using mMessage Machine T7 Transcription Kit (Thermo Fisher Scientific). RNA was purified using RNEasy kit (Qiagen) and stored as aliquoted stocks at -80 °C. Defolliculated stage V-VI oocytes were microinjected with 11-13 ng mRNA and incubated at 18°C for expression. Currents were recorded within 48-54 h after microinjection. For amplitude comparisons, currents were compared at the same time of expression for sample injected with equivalent amounts of mRNA.

For recordings, oocytes were impaled with borosilicate recording microelectrodes (0.2-–2.0 MΩ resistance) backfilled with 3 M KCl and were subjected to constant perfusion of ND96 at a rate of 3-5 ml min^−1^. Except where otherwise indicated, recording solution was 96 mM NaCl, 2 mM KCl, 1.8 mM CaCl_2_, and 1.0 mM MgCl_2_, buffered with 5 mM HEPES, pH 7.4. For each recording, control solution (ND96) was perfused over a single oocyte until the current stabilized. Currents were evoked from a -80 mV holding potential followed by a 500 ms ramp from -140 mV to +50 mV. Data were recorded using a Axoclamp 900A amplifier (Molecular Devices) controlled by pCLAMP10.7 software (Molecular Devices), and digitized at 1 kHz using Digidata 1550B digitizer (Molecular Devices). Representative traces and dose response plots were generated in GraphPad Prism 10 (GraphPad Software, Boston).

#### Data quantification and statistical analysis

The results are from 2-4 independent oocyte batches. *P* values for Fig. 4c are as follows: *P* < 0.9999 (K_2P_6.1(Twik2) vs uninjected oocytes), *P* < 0.0001 (TWIK-2 vs TWIK-2_RM_), *P* < 0.0001 (TWIK-2_RM_ vs TWIK-2_RM_ V131D), *P* = 0.9768 (TWIK-2_RM_ vs TWIK-2_RM_ M135A), *P* < 0.0001 (TWIK-2_RM_ vs TWIK-2_RM_ M135D), *P* < 0.0001 (TWIK-2_RM_ vs TWIK-2_RM_ R257A) and *P* < 0.0001 (TWIK-2_RM_ vs TWIK-2_RM_ R257E). Statistical analysis was performed using One-Way ANOVA and multiple comparisions are performed with Tukey’s test. Error bars are S.E.M.. Full detailed statistical information is available in the source data spreadsheet.

*P* values for Fig. 4e are as follows: *P* < 0. 9999 (TWIK2 vs uninjected oocytes), *P* < 0.0001 (TWIK-2 vs TWIK-2_RM_), *P* = 0.0341 (TWIK-2_RM_ vs TWIK-2_RM_ Y111A), *P* = 0. 9999 (TWIK-2_RM_ vs TWIK-2_RM_ Y111H) and *P* = 0.0068 (TWIK-2_RM_ vs TWIK-2_RM_ Y111N). Statistical analysis was performed using One-Way ANOVA and multiple comparisions are performed with Tukey’s test. Error bars are S.E.M.. Full detailed statistical information is available in the source data spreadsheet.

### Data Availability

K_2P_6.1 (TWIK2) nanodisc and detergent structures (PDB: 9OTA, EMDB-70828 and 9OTK, EMDB-70837, respectively), K_2P_6.1 (TWIK2)_RM_ plugged and unplugged structures (PDB: 9OV0, EMDB-70885 and 9OV9, EMDB-70894, respectively), and K_2P6.1_ (TWIK2)_RM_ R257A plugged and unplugged structures (PDB:9OVD, EMDB-70898 and 9OVE, EMDB-70899, respectively). Human K_2P_6.1 (TWIK2) sequence is Uniprot Q9Y257.

Data or materials will be provided on request from the corresponding author.

## Notes

### Competing Interest Statement

The authors have declared no competing interest.

