## Supplementary material for "A lipid plug affects K_2P_6.1(TWIK-2) function": Supplmentary Figures S1-S18

11 June 2025

[6]Molecular Biophysics and Integrated Bio-imaging Division
Lawrence Berkeley National Laboratory, Berkeley, CA 94720 USA

**Figure S1** **Mondal, et al.**

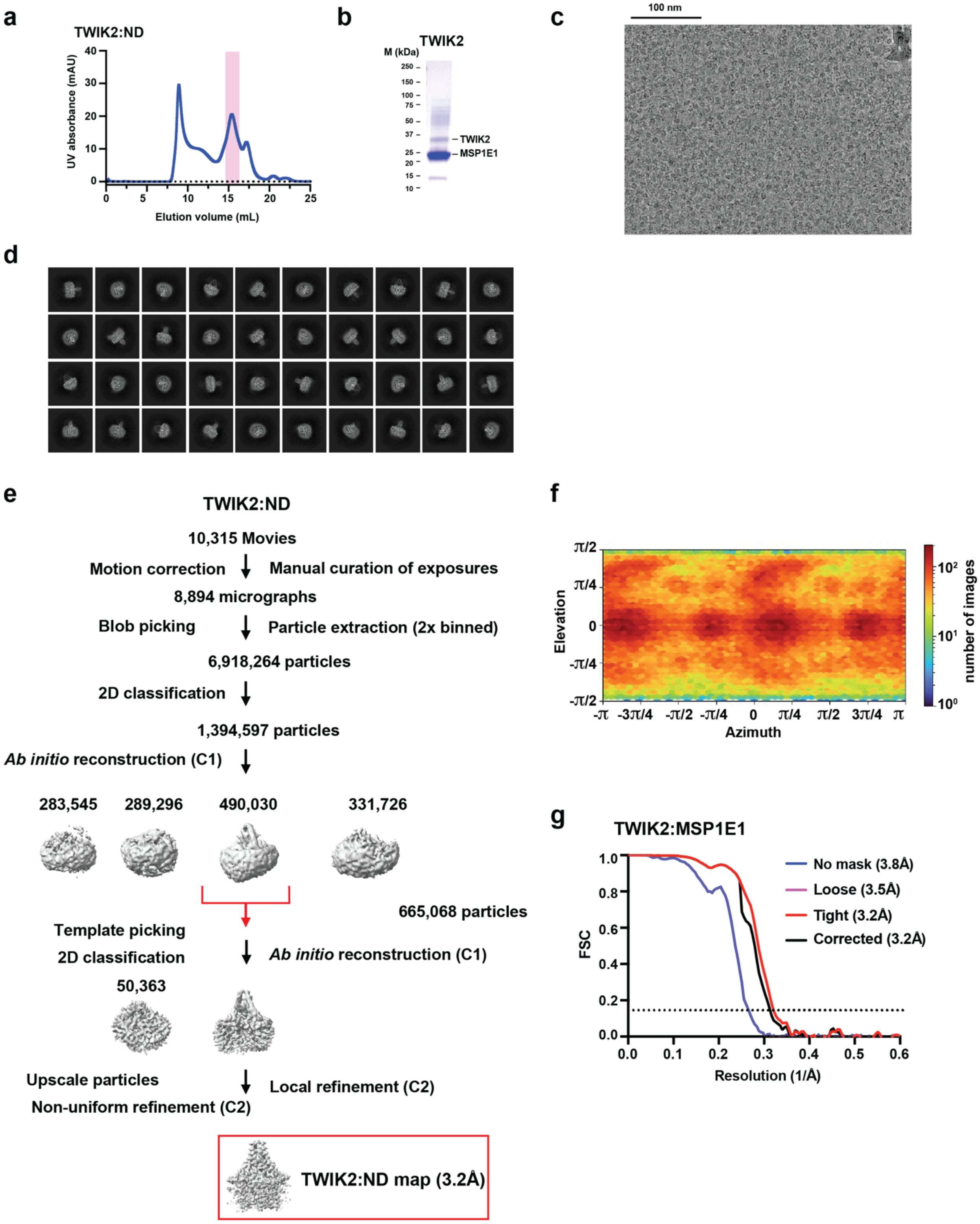

**Figure S1 Cryo-EM analysis of TWIK2 in MSP1E1 nanodiscs** Exemplar **a,** SEC (Superose 6 Increase 10/300 GL) for TWIK2:MSP1E1 nanodiscs (TWIK2:ND), **b,** peak fraction SDS-PAGE. **c,** electromicrograph (~105,000x magnification), and **d,** 2D class averages. **e,** Workflow for electron microscopy data processing for the TWIK2:MSP1E1 nanodisc complex in cryoSPARC-3.2[1]. Red arrow indicates the class of particles extracted without Fourier cropping after the initial cleanup. These were further subjected to heterogeneous, non-uniform, and local refinement jobs that resulted in the final map at 3.2Å (red box). **f,** Particle distribution plot and **g,** gold-standard Fourier Shell Correlation (FSC) curves for the TWIK2:MSP1E1 nanodisc complex.

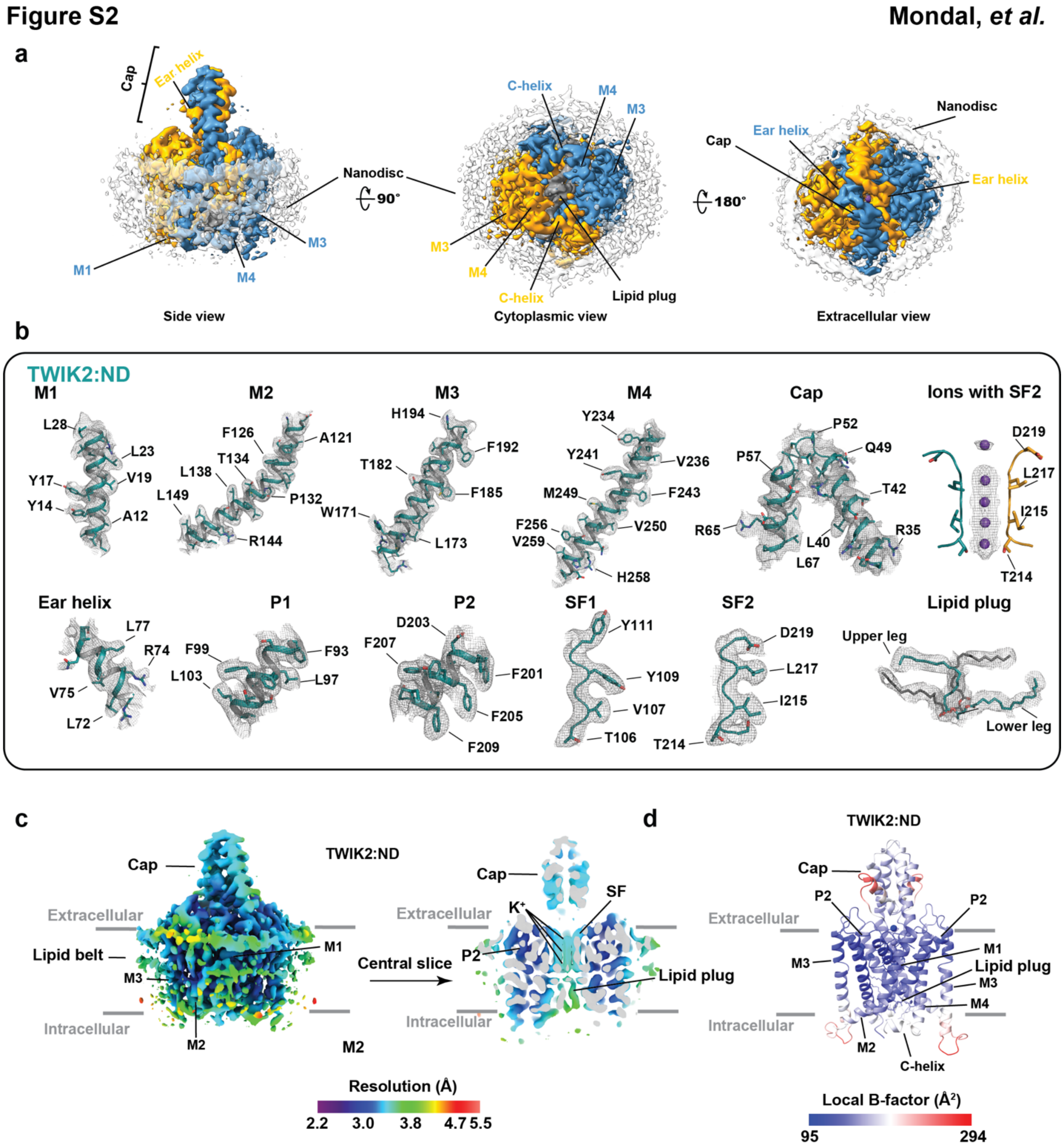


**Figure S2 TWIK2:ND cryo-EM map and model quality a,** CryoEM map of the TWIK2:MSP1E1 complex. TWIK2 subunits are colored blue and yellow. Lipid plug is colored grey. Nanodisc is clear. Locations of select channel elements are indicated. **b,** Electron microscopy maps for indicated TWIK2 elements. Select residues are indicated. Channel elements are deep teal and bright orange. Potassium ions are purple. Lipid plug is deep teal and grey. Maps are rendered at 2-4σ. **c,** TWIK2:ND local resolution showing a central slice through the channel. Select channel elements and lipid belt s are labeled. **d,** TWIK2:ND local B-factor.

**Figure S3** **Mondal, *et al.***

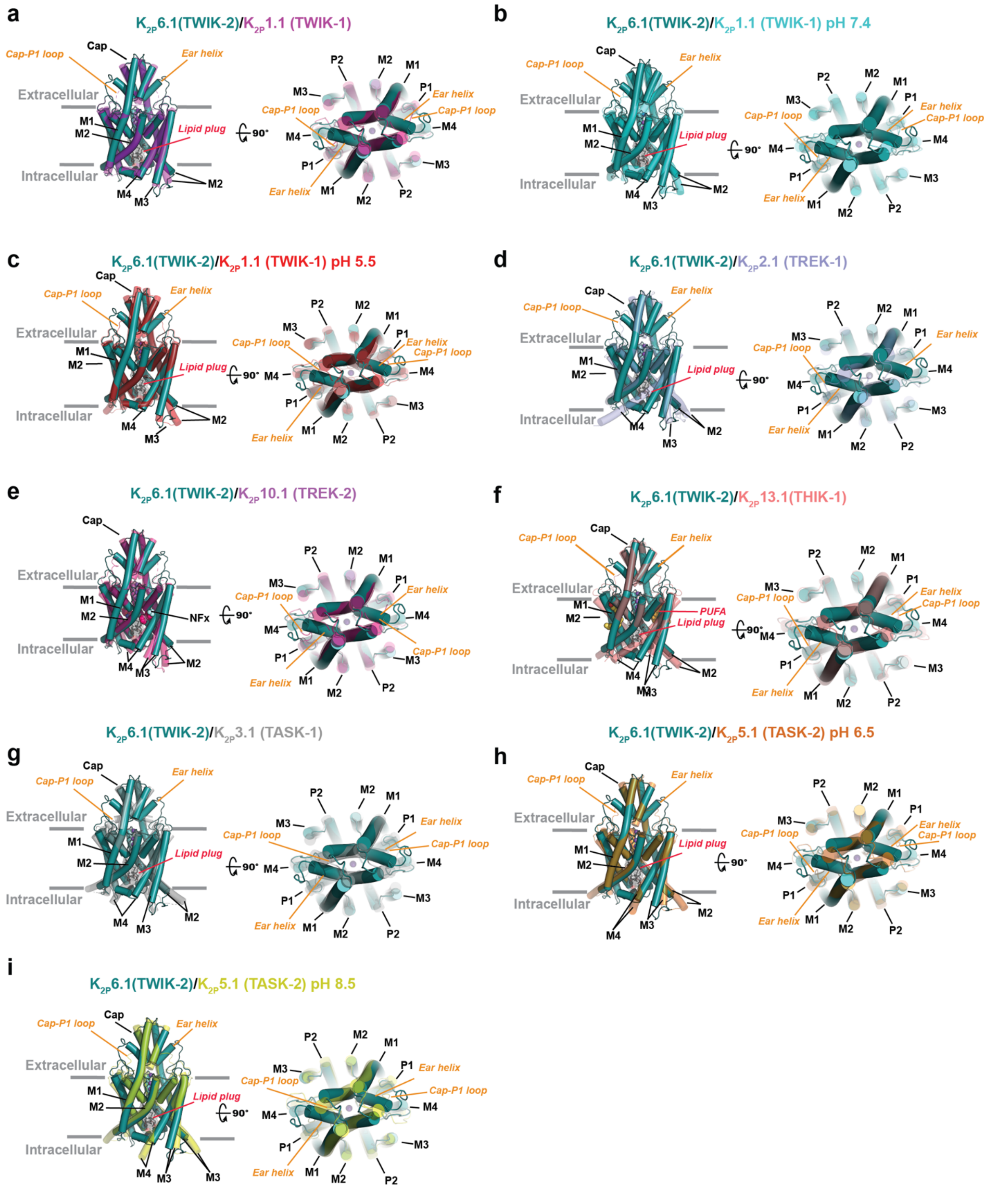


**Figure S3 K2P6.1 (TWIK-2) structure comparisons.** Superpositions of K2P6.1 (TWIK-2) (deep teal) with: **a,** K2P1.1 (TWIK-1) (magenta) (PDB:3UKM)[2] RMSD$_{C\alpha}$ = 0.817 Å); **b,** K2P1.1 (TWIK-

1) pH 7.4 (cyan) (PDB:7SK0) [3](RMSD$_{C\alpha}$ = 1.016 Å) **c,** $K_{2P}$1.1 (TWIK-1) pH 5.5 (red) (PDB:7SK1) [3] (RMSD$_{C\alpha}$ = 1.089 Å); **d,** $K_{2P}$2.1 (TREK-1) (light blue) (PDB: 6CQ6) [4]( RMSD$_{C\alpha}$ = 1.021 Å); **e,** $K_{2P}$10.1(TREK-2):norfluoxetine (Nfx) complex (hot pink) (PDB:4XDK) (RMSD$_{C\alpha}$ = 1.05 Å) [5]; **f,** $K_{2P}$13.1(THIK-1) (deep salmon) (PDB:9BSN) [6]( RMSD$_{C\alpha}$ = 0.996); **g,** $K_{2P}$3.1(TASK-1) (grey) (PDB:6RV2) (RMSD$_{C\alpha}$ = 1.051 Å); **h,** $K_{2P}$5.1(TASK-2) pH 6.5 (orange) (PDB:6WLV) [7] (RMSD$_{C\alpha}$ = 1.071); and **i,** $K_{2P}$5.1(TASK-2) pH 8.5 (yellow) (PDB:6WM0) [7] (RMSD$_{C\alpha}$ = 0.978). Each panel shows side (left) and extracellular (right) views. Grey bars denote the membrane.

**Figure S4** **Mondal, *et al.***

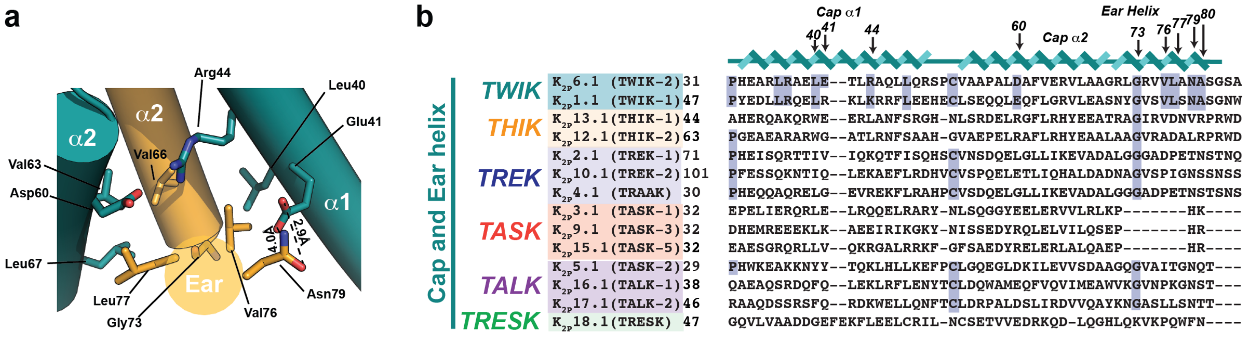


**Figure S4 K2P6.1 (TWIK-2) Ear helix interactions and sequence comparison. a,** Local environment showing K2P6.1 (TWIK-2) Ear helix-Cap interactions. K2P6.1 (TWIK-2) subunits (deep teal and bright orange) and select residues are indicated. Dashes lines indicate hydrogen bond interactions. **b,** Sequence comparison of Cap and Ear helix from K2P6.1 (TWIK-2) with other K2PS. Numbers indicate residues shown in 'a'. Conservation is shown in blue. Sequences in 'b' are for human: K2P6.1(TWIK-2) (GENBANK 4758624); K2P 1.1(TWIK-1) (GENBANK 4504847); K2P13.1(THIK-1) (GENBANK 16306555); K2P12.1(THIK-2) (GENBANK 11545761); K2P2.1(TREK-1) (GENBANK 14589851); K2P10.1(TREK-2) (GENBANK 20143944); K2P4.1(TRAAK) (GENBANK 15718767); K2P3.1(TASK-1) (GENBANK 4504849; K2P9.1 (TASK-3) (GENBANK 542133161); K2P15.1(TASK-5) (GENBANK 333440483); K2P 5.1(TASK-2) (GENBANK 4504851); K2P16.1(TALK-1) (GENBANK 14149764); K2P17.1(TALK-2) (GENBANK 17025230); and K2P18.1(TRESK) (GENBANK 32469495).

**Figure S5** **Mondal, *et al.***

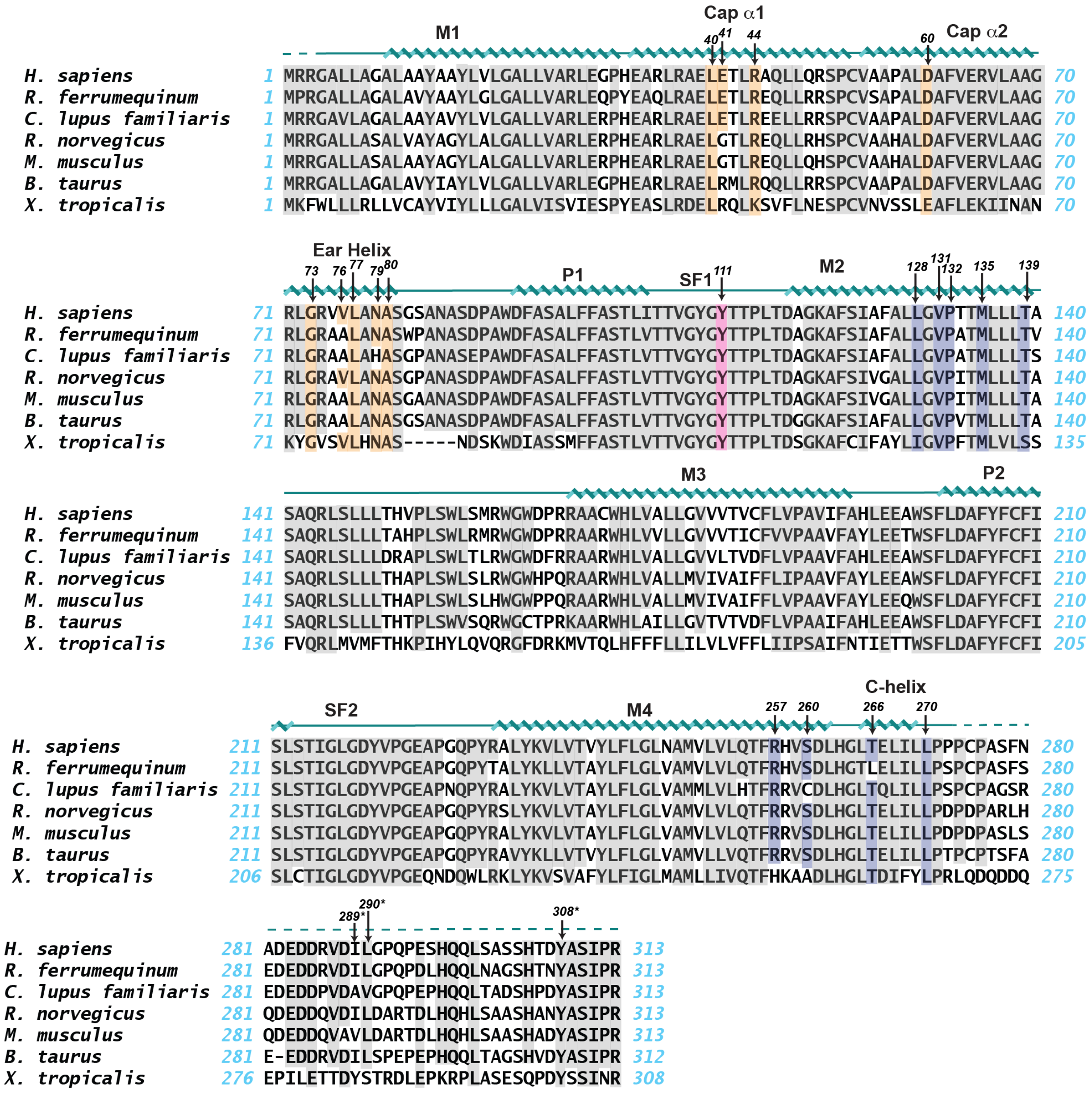


**Figure S5 $K_{2P}6.1$ (TWIK-2) sequence conservation.** Sequence alignment of *H. sapiens* $K_{2P}6.1$(TWIK-2) (GENBANK 4758624**)** with homologs from the greater horseshoe bat (*R. ferrumequinum*) (GENBANK 117034693), domestic dog (*C. lupus familiaris*) (GENBANK 484531), rat (*R. norvegicus*) (GENBANK 116491), mouse (*M. musculus*) (GENBANK 75766694), cattle (B. taurus) (GENBANK 785418), and Western clawed frog (*X. tropicalis*) (GENBANK 100145504). Cap and Ear helix interaction sites (orange), Tyr111 (magenta), and plug lipid contact sites (blue) are indicated. Asterisk indicates sites of I289A/L290A/Y308 retention mutations. Dashed lines indicate residues lacking defined density in the TWIK2:ND structure.

**Figure S6** **Mondal, et al.**

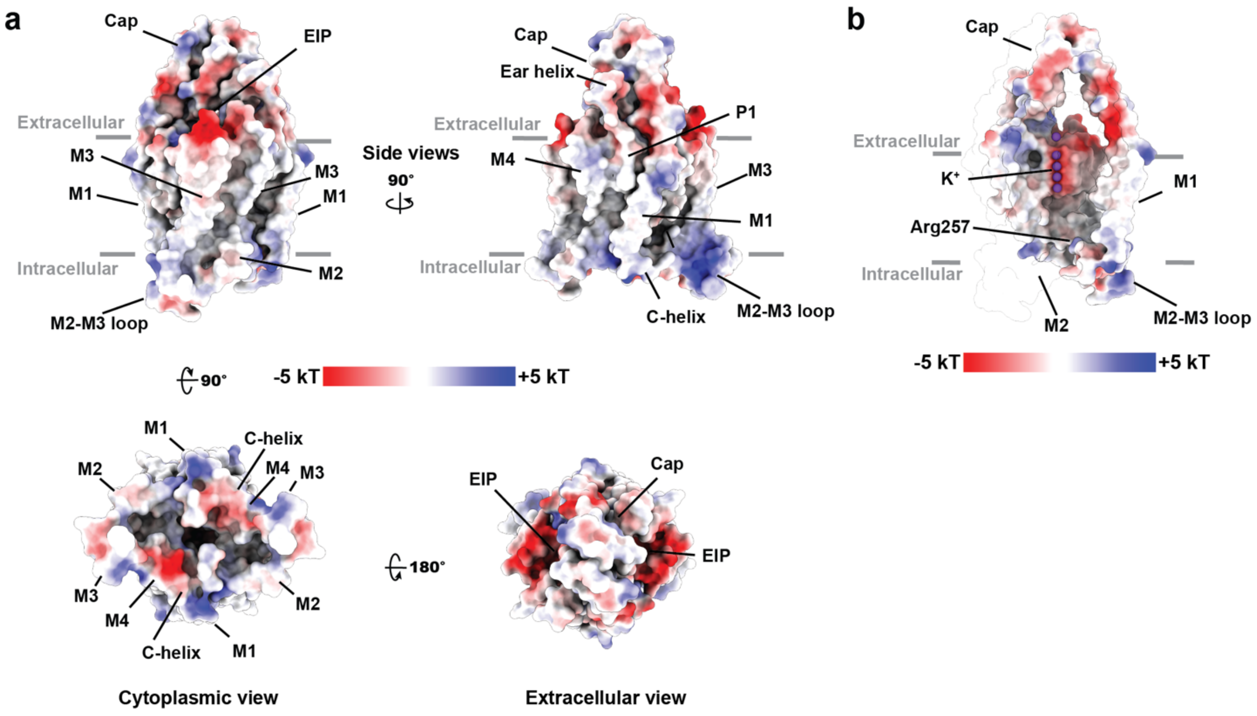


**Figure S6 K2P6.1 (TWIK-2) electrostatic surface potentials. a,** Electrostatic surface potentials calculated using APBS [8]. **b,** Slice through the center of K2P6.1 (TWIK-2) showing the inner cavity. Selectivity filter ions are shown as purple spheres. Select channel elements are labeled. Grey bars denote the membrane.

**Figure S7** **Mondal, et al.**

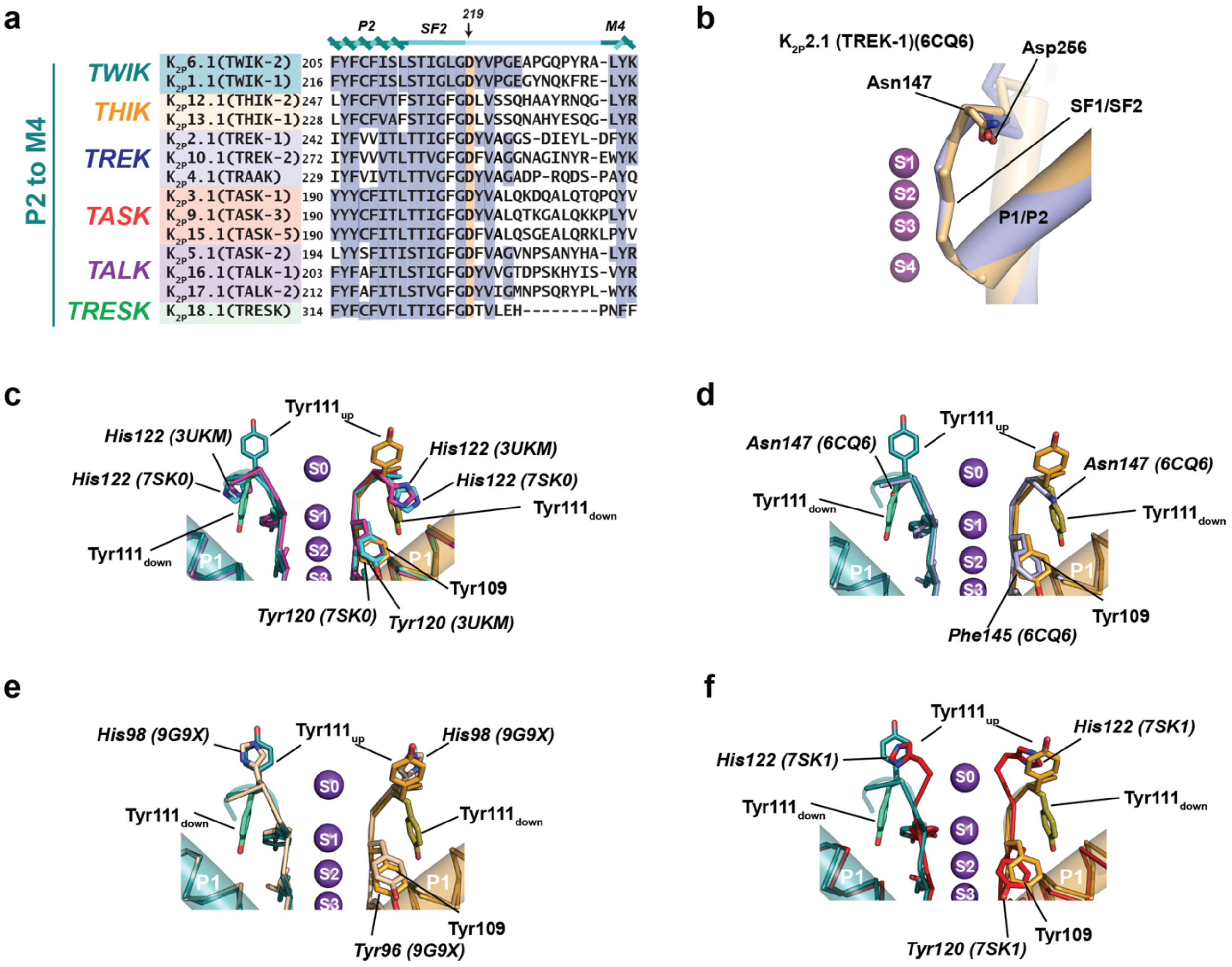


**Figure S7 K2P6.1 (TWIK-2) selectivity filter comparisons. a,** Sequence comparison of P2 through initial segment of M4 from K2P6.1 (TWIK-2) with other K2Ps. Site corresponding to Y111 in SF1 is labeled and indicated in orange. Conservation is shown in blue. SF2-M4 linker is denoted in light blue. **b,** Superposition of SF1 (light orange) and SF2 (slate) regions from K2P2.1(TREK-1) (PDB: 6CQ6) [4] showing the shared conformation of the Asn147 and Asp226 sites corresponding to K2P6.1 (TWIK-2) Tyr111. **c-f,** Superposition of P1-SF1 from K2P6.1 (TWIK-2) with **c,** K2P1.1(TWIK-1) (magenta) (PDB:3UKM)[2] and K2P1.1 (TWIK-1) pH 7.4 (cyan) (PDB:7SK0) [3]; **d,** K2P2.1(TREK-1) (PDB: 6CQ6) [4]; **e,** K2P3.1 (TASK-1) (wheat) pH 7.5 (PDB:9G9X) [9]; **f,** K2P1.1(TWIK-1) pH 5.5 (red) (PDB:7SK1) [3]. In 'c-f' the Tyr111 'up' and 'down' conformations are indicated. Sequences in 'a' are for human: K2P6.1(TWIK-2) (GENBANK 4758624); K2P 1.1(TWIK-1) (GENBANK 4504847); K2P13.1(THIK-1) (GENBANK 16306555);

$K_{2P}$12.1(THIK-2) (GENBANK 11545761); $K_{2P}$2.1(TREK-1) (GENBANK 14589851); $K_{2P}$10.1(TREK-2) (GENBANK 20143944); $K_{2P}$4.1(TRAAK) (GENBANK 15718767); $K_{2P}$3.1(TASK-1) (GENBANK 4504849); $K_{2P}$9.1 (TASK-3) (GENBANK 542133161); $K_{2P}$15.1(TASK-5) (GENBANK 333440483); $K_{2P}$ 5.1(TASK-2) (GENBANK 4504851); $K_{2P}$16.1(TALK-1) (GENBANK 14149764); $K_{2P}$17.1(TALK-2) (GENBANK 17025230); and $K_{2P}$18.1(TRESK) (GENBANK 32469495).

### Figure S8

Mondal, *et al.*

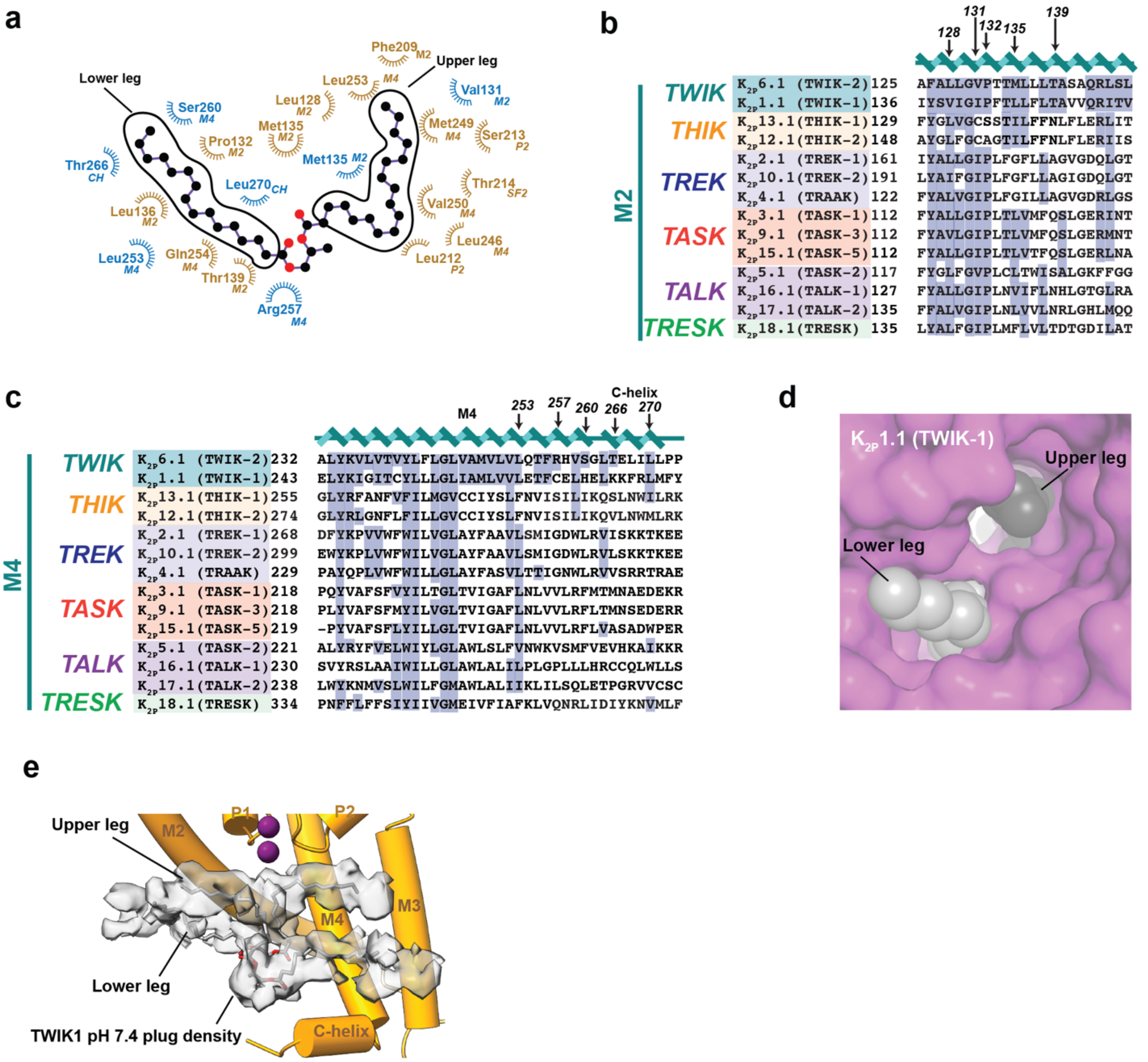


**Figure S8 $K_{2P}$6.1 (TWIK-2):plug lipid interactions. a,** Ligplot[10] diagram showing plug lipid interactions (≤ 5Å). $K_{2P}$6.1 (TWIK-2) chains are indicated in blue and olive. Lipid upper and lower leg elements are indicated. Channel elements M2, M4, selectivity filter 2 (SF2), and C-helix (CH) are indicated. **b,** and **c,** Sequence comparison of $K_{2P}$6.1 (TWIK-2) **b,** M2 and **c,** M4 and C-helix regions with other with other $K_{2P}$s. Labels indicated residues that interact with the plug lipid. Conservation is indicated in blue. **d,** View of the $K_{2P}$1.1(TWIK-1) (PDB:3UKM) [2] surface from the perspective of the bilayer. Lipids (grey and black) are from the TWIK2 structure. View is similar to Fig. 2d. Sequences in 'b' and 'c' are for human: $K_{2P}$6.1(TWIK-2) (GENBANK 4758624); $K_{2P}$ 1.1(TWIK-1) (GENBANK 4504847); $K_{2P}$13.1(THIK-1) (GENBANK 16306555);

$K_{2P}$12.1(THIK-2) (GENBANK 11545761); $K_{2P}$2.1(TREK-1) (GENBANK 14589851); $K_{2P}$10.1(TREK-2) (GENBANK 20143944); $K_{2P}$4.1(TRAAK) (GENBANK 15718767); $K_{2P}$3.1(TASK-1) (GENBANK 4504849); $K_{2P}$9.1 (TASK-3) (GENBANK 542133161); $K_{2P}$15.1(TASK-5) (GENBANK 333440483); $K_{2P}$5.1(TASK-2) (GENBANK 333440483); $K_{2P}$16.1(TALK-1) (GENBANK 14149764); $K_{2P}$17.1(TALK-2) (GENBANK 17025230); and $K_{2P}$18.1(TRESK) (GENBANK 32469495). **e,** Lipid plug density (1.8$\sigma$) from $K_{2P}$1.1 (TWIK-1) at pH 7.4 (EMDB 25168)[3]. Model shows a single $K_{2P}$6.1(TWIK-2) subunit and lipid plug (grey)

**Figure S9** **Mondal, et al.**

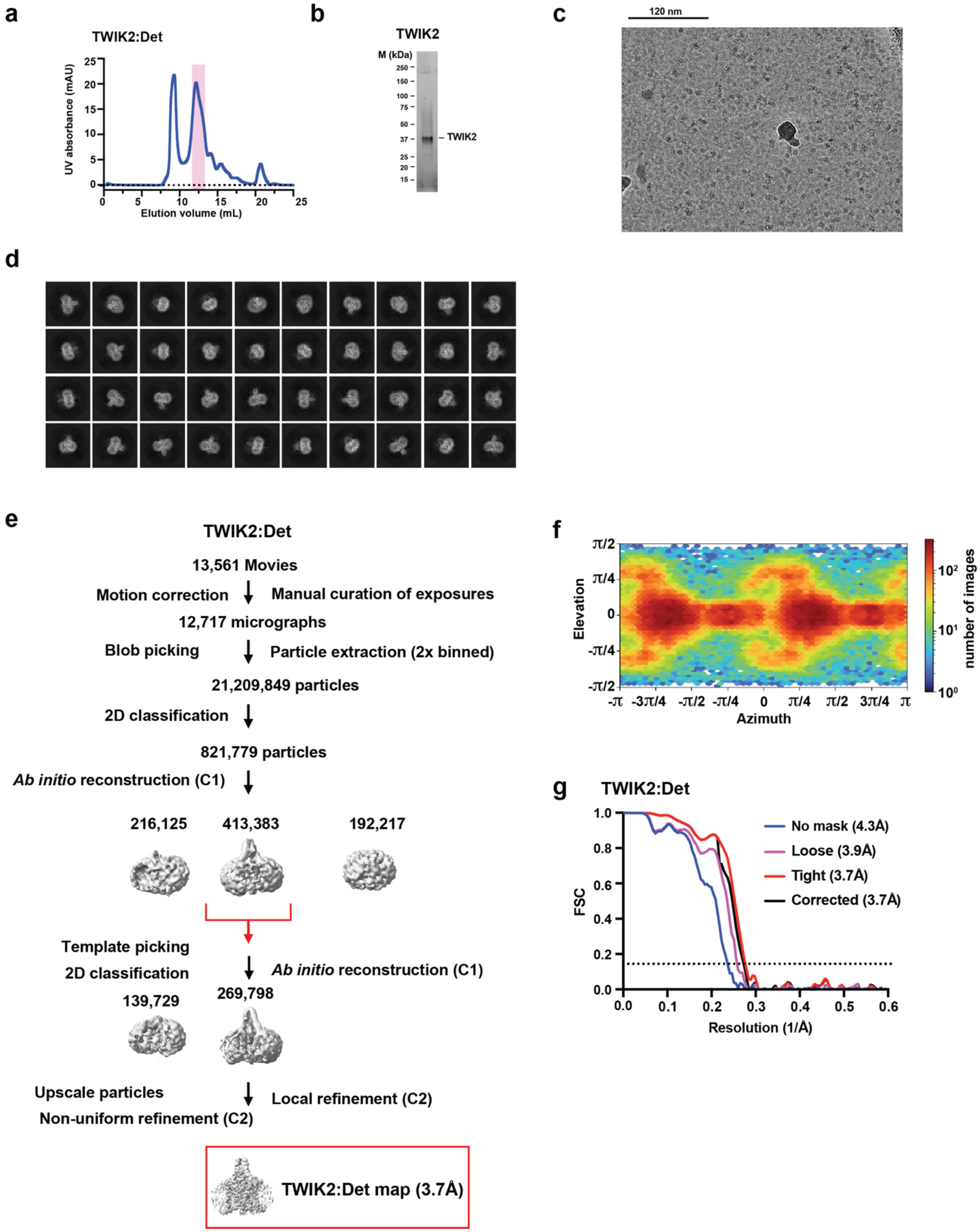

**Figure S9 Cryo-EM analysis of TWIK2 in detergent.** Exemplar **a,** SEC (Superose 6 Increase 10/300 GL) for TWIK2 in β-docecylmaltoside (TWIK2:Det), **b,** peak fraction SDS-PAGE. **c,** electromicrograph (~105,000x magnification), and **d**, 2D class averages. **e,** Workflow for electron microscopy data processing for TWIK2:Det in cryoSPARC-3.2[1]. Red arrow indicates the class of particles extracted without Fourier cropping after the initial cleanup. These were further subjected to heterogeneous, non-uniform, and local refinement jobs that resulted in the final map at 3.7Å (red box). **f,** Particle distribution plot and **g,** gold-standard Fourier Shell Correlation (FSC) curves for TWIK2:Det.

Figure S10 Mondal, *et al.*

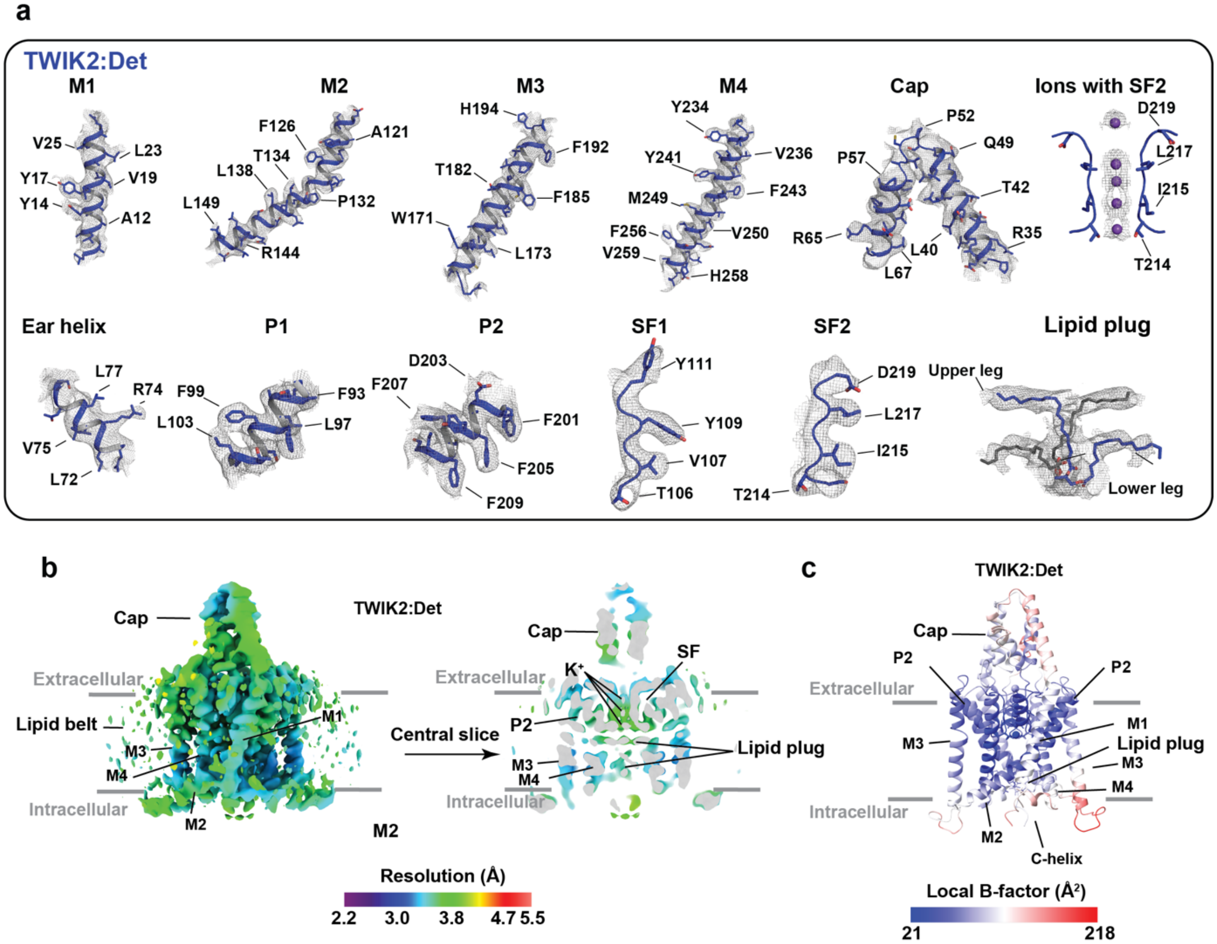


**Figure S10 TWIK2:Det cryo-EM map and model quality a**, Electron microscopy maps for indicated TWIK2 elements. Select residues are indicated. Channel elements are blue. Potassium ions are purple. Lipid plug is blue and grey. Maps are rendered at 3-4σ. **c,** TWIK2:Det local resolution showing a central slice through the channel. Select channel elements and lipid belt s are labeled. **d,** TWIK2:Det local B-factor.

**Figure S11** **Mondal, et al.**

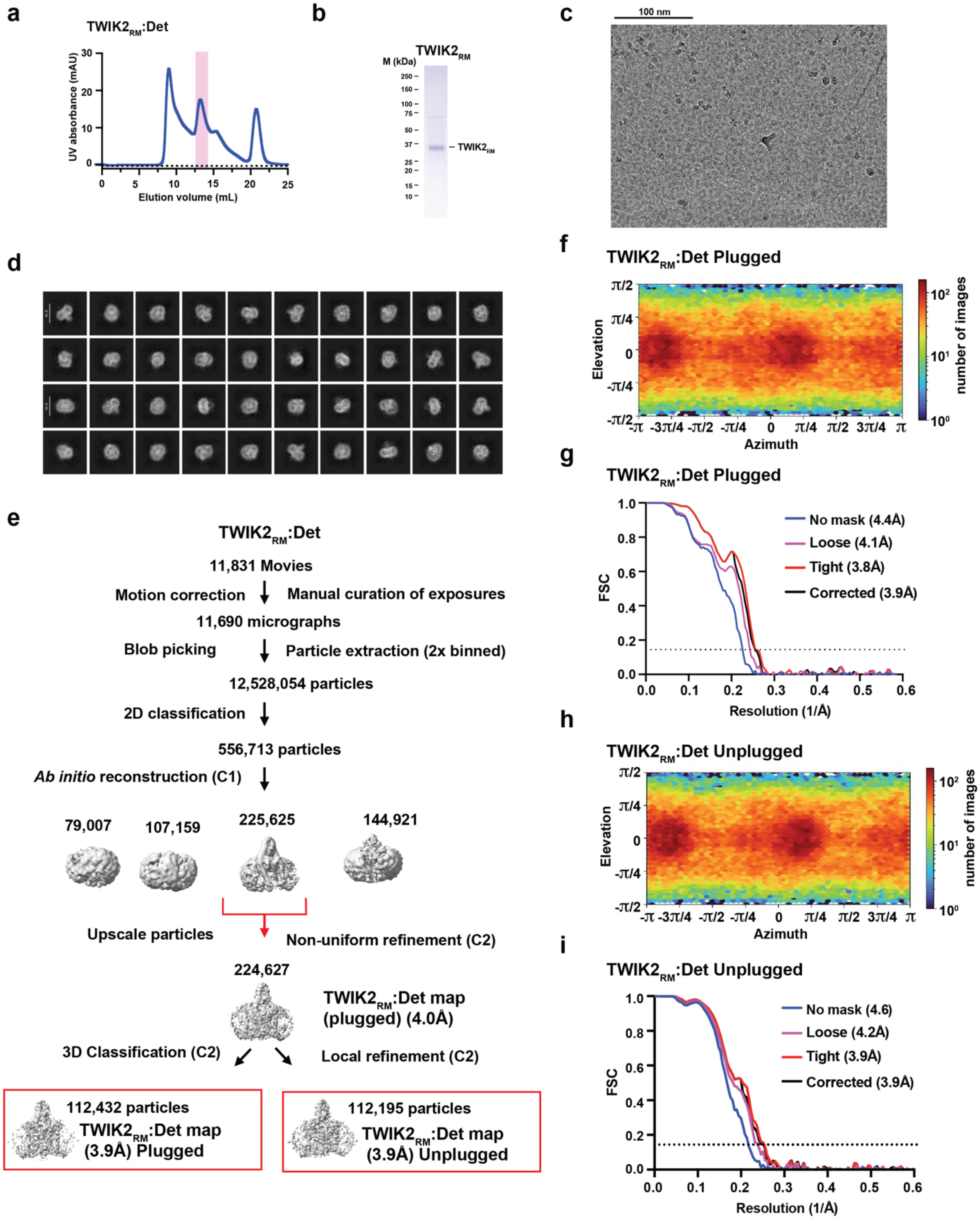


**Figure S11 Cryo-EM analysis of $TWIK2_{RM}$ in detergent.** Exemplar **a,** SEC (Superose 6 Increase 10/300 GL) for $TWIK2_{RM}$ in N-dodecyl-β–D-maltoside (TWIK2:Det), **b,** peak fraction

SDS-PAGE. **c,** electromicrograph (~105,000x magnification), and **d**, 2D class averages. **e,** Workflow for electron microscopy data processing for $TWIK2_{RM}$:Det in cryoSPARC-3.2[1]. Red arrow indicates the class of particles extracted without Fourier cropping after the initial cleanup. Non-uniform refinement, followed by 3D classification was performed using the extracted particles yielding 6 different classes that were then combined, following manual inspection, into two classes representing the plugged and unplugged channel. These particle-sets were further subjected to Local Refinement to obtain the final maps. **f,** Particle distribution plot and **g,** gold-standard Fourier Shell Correlation (FSC) curves for $TWIK2_{RM}$ Plugged. **h,** Particle distribution plot and **i,** gold-standard Fourier Shell Correlation (FSC) curves for $TWIK2_{RM}$ Unplugged.

**Figure S12** **Mondal, *et al.***

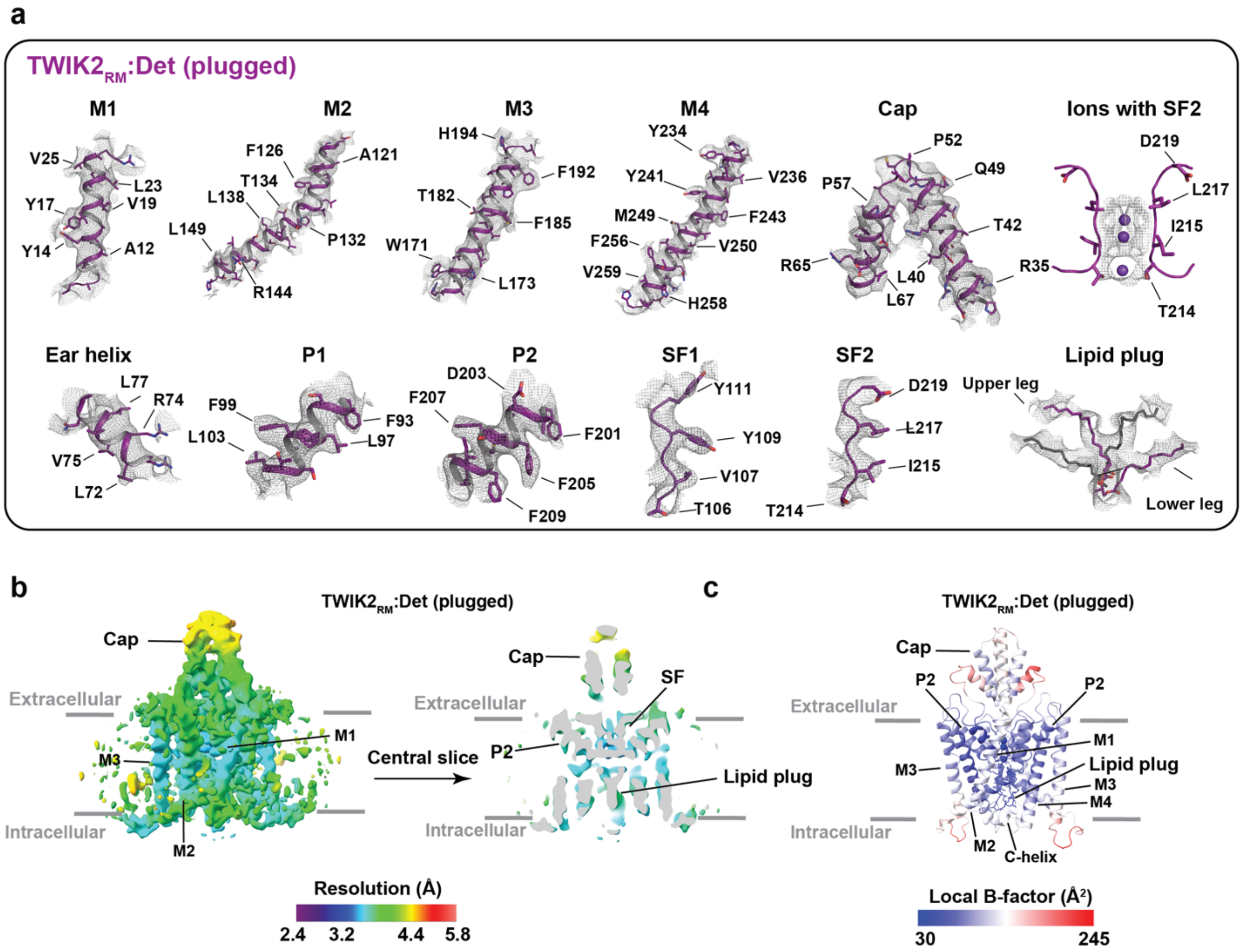


**Figure S12 TWIK2$_{RM}$:Det (plugged) cryo-EM map and model quality a**, Electron microscopy maps for indicated TWIK2$_{RM}$ (plugged) elements. Select residues are indicated. Channel elements are dark purple. Potassium ions are purple. Lipid plug is dark purple and grey. Maps are rendered at 3-4σ. **c,** TWIK2:Det$_{RM}$ (plugged) local resolution showing a central slice through the channel. Select channel elements and lipid belt s are labeled. **d,** TWIK2:Det$_{RM}$ (plugged) local B-factor.

**Figure S13** **Mondal, *et al.***

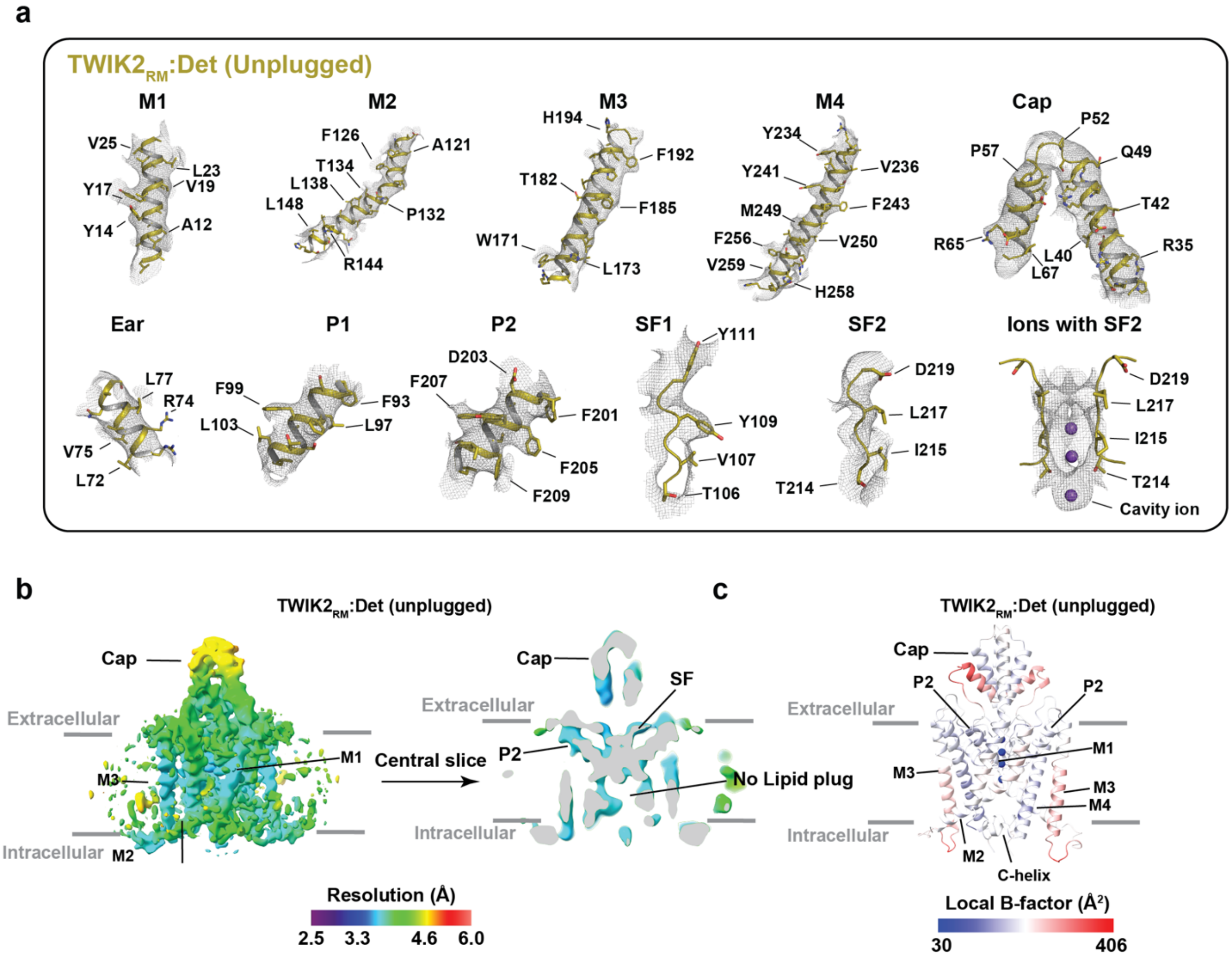


**Figure S13 TWIK2RM:Det (unplugged) cryo-EM map and model quality a**, Electron microscopy maps for indicated TWIK2RM (unplugged)elements. Select residues are indicated. Channel elements are dark purple. Potassium ions are purple. Lipid plug is dark purple and grey. Maps are rendered at 3-4σ. **c,** TWIK2RM:Det (unplugged) local resolution showing a central slice through the channel. Select channel elements are labeled. **d,** TWIK2:DetRM (unplugged) local B-factor.

**Figure S14** **Mondal, *et al.***

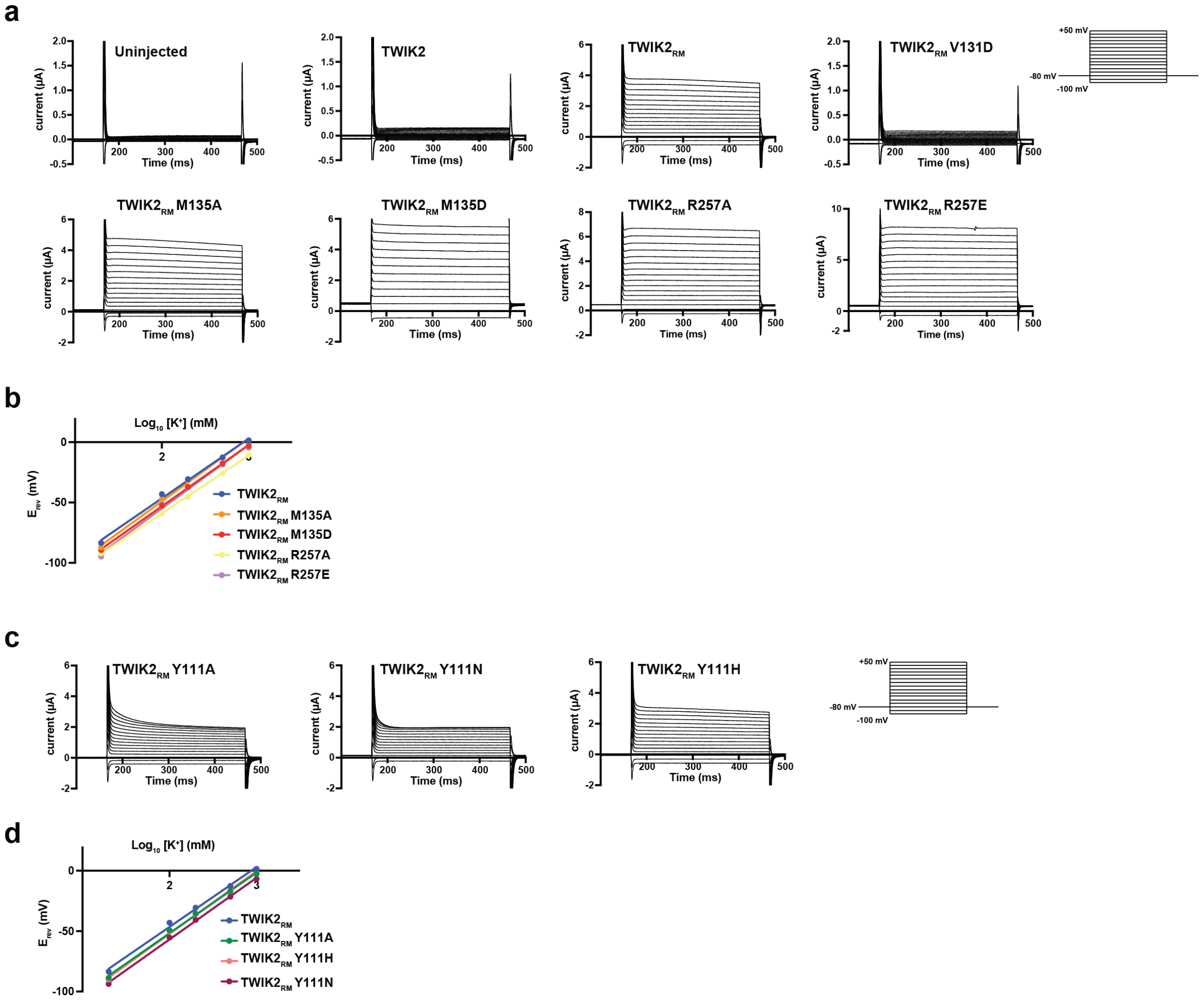


**Figure S14 Functional studies of TWIK2 and mutants. a,** Exemplar TEVC recordings from *Xenopus* oocytes. Currents for uninjected (control), TWIK2, TWIK2$_{RM}$ and TWIK2$_{RM}$ mutants V131D, M135A, M135D, R257A, and R257E. Inset shows protocol. **b,** Potassium selectivity for TWIK2$_{RM}$ (blue) TWIK2$_{RM}$ M135A (orange), TWIK2$_{RM}$ M135D (red), TWIK2$_{RM}$ R257A (yellow), and TWIK2$_{RM}$ R257E (purple). **c,** Exemplar TEVC recordings from *Xenopus* oocytes. Currents for TWIK2$_{RM}$ Y111A, TWIK2$_{RM}$ Y111N, and TWIK2$_{RM}$ Y111H. Inset shows protocol. **d,** Potassium selectivity for TWIK2$_{RM}$ (blue) TWIK2$_{RM}$ Y111A (green), TWIK2$_{RM}$ Y111H (orange), TWIK2$_{RM}$ Y111N (maroon). In 'b' and 'd' lines represent fits to the Nernst equation $E_K = RT/zF \ln[K^+]_{out}/[K^+]_{in}$ assuming $/[K^+]_{in} = 108.6$ mM [11].

**Figure S15** **Mondal, et al.**

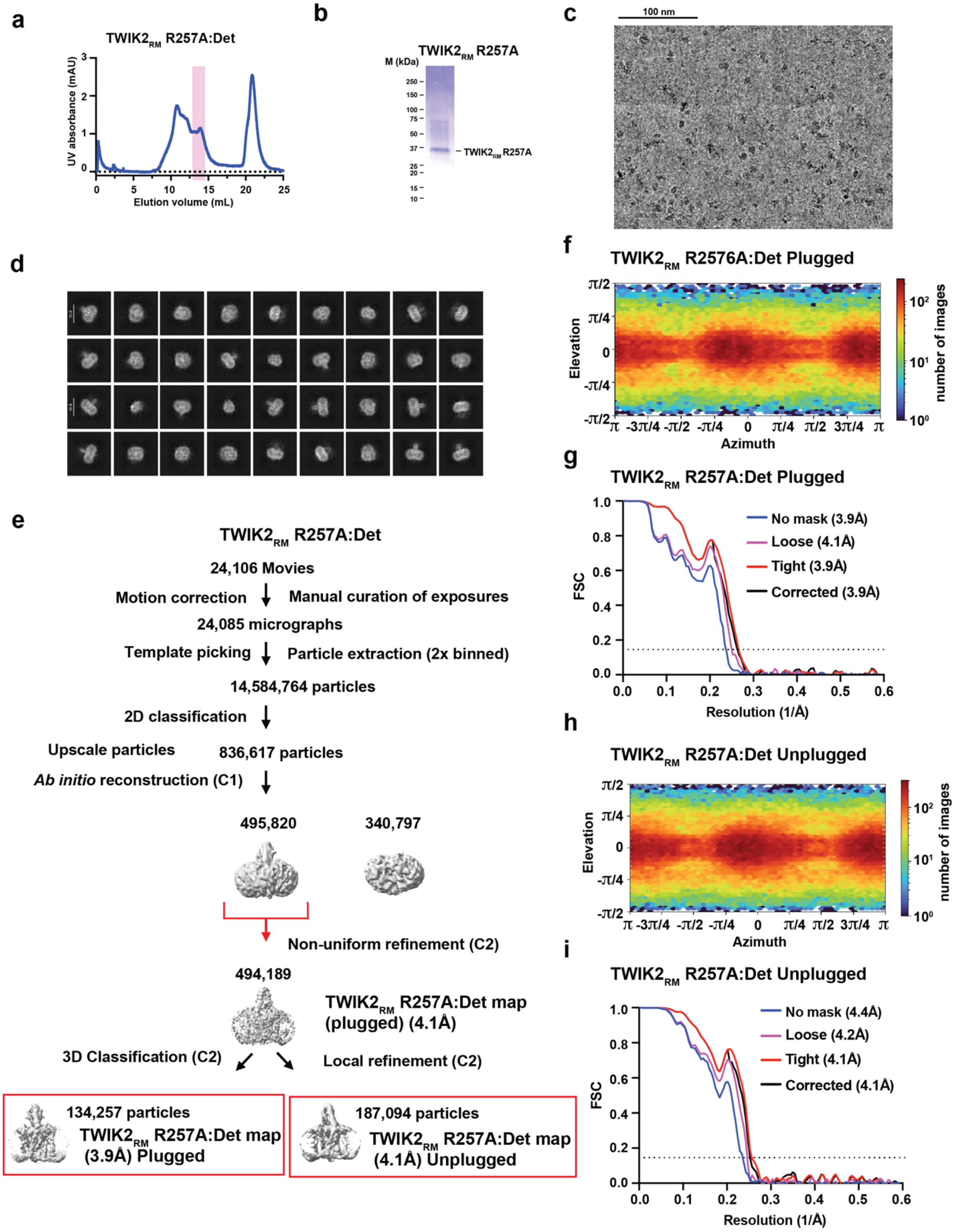

**Figure S15 Cryo-EM analysis of TWIK2$_{RM}$ R257A in detergent.** Exemplar **a,** SEC (Superose 6 Increase 10/300 GL) for TWIK2$_{RM}$ R257A in N-dodecyl-β–D-maltoside (TWIK2:Det), **b,** peak fraction SDS-PAGE. **c,** electron micrograph (~105,000x magnification), and **d**, 2D class averages. **e,** Workflow for electron microscopy data processing for TWIK2$_{RM}$ R257A:Det in cryoSPARC-3.2[1]. Red arrow indicates the class of particles re-extracted without Fourier cropping after the initial cleanup. Extracted particles were further subjected to non-uniform refinement to generate a consensus map. This refined particle set was used for 3D classification having lipid plug density as focused mask and generated 6 classes, which later combined into two major groups having either the presence or absence of the lipid plug. These particle-sets were further subjected to Local Refinement to obtain the final maps. **f,** Particle distribution plot and **g,** gold-standard Fourier Shell Correlation (FSC) curves for TWIK2$_{RM}$ R257A Plugged. **h,** Particle distribution plot and **i,** gold-standard Fourier Shell Correlation (FSC) curves for TWIK2$_{RM}$ R257A Unplugged.

**Figure S16** **Mondal, *et al.***

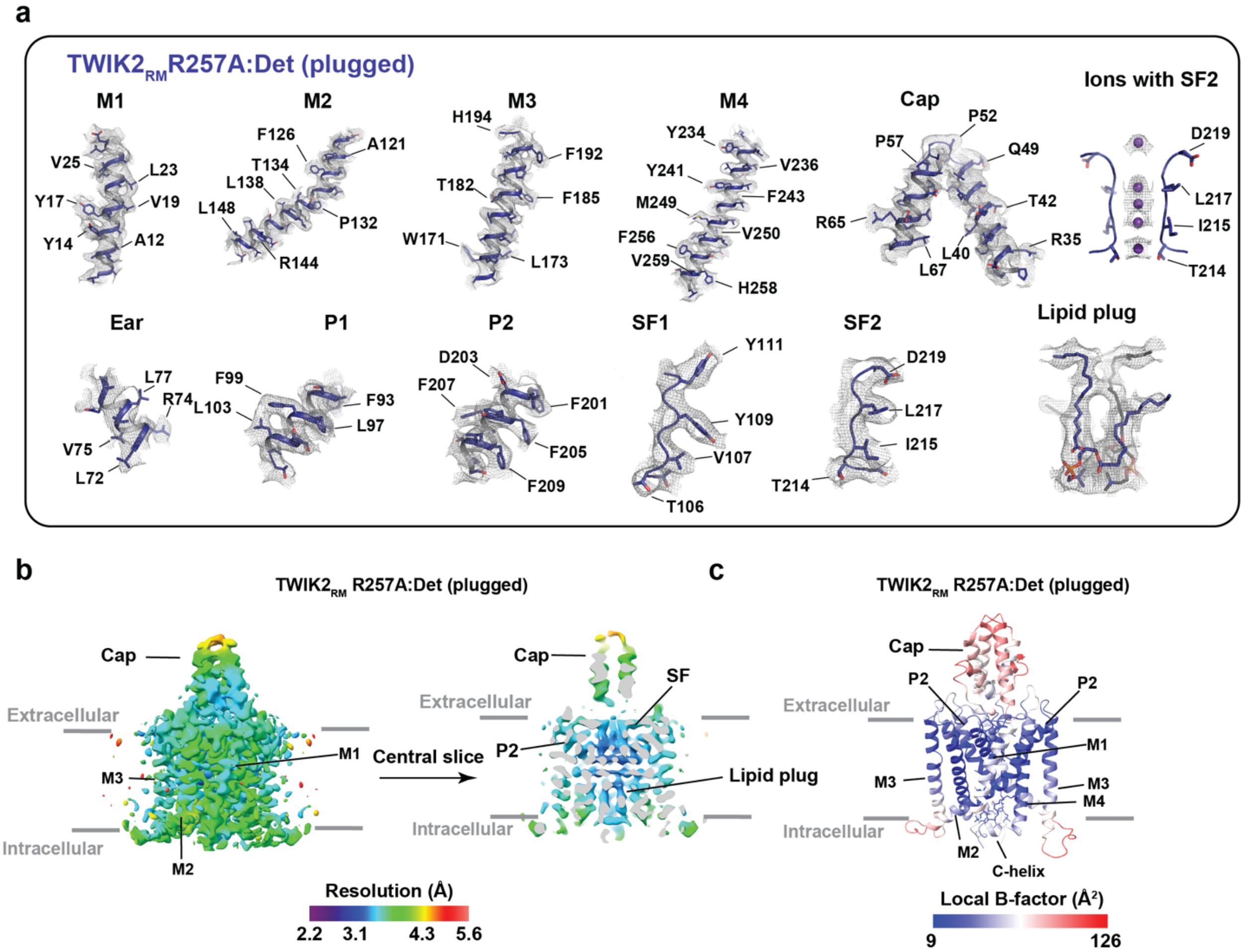


**Figure S16 TWIK2RM R257A:Det (plugged) cryo-EM map and model quality a**, Electron microscopy maps for indicated TWIK2RM R257A:Det (plugged) elements. Select residues are indicated. Channel elements are dark purple. Potassium ions are purple. Lipid plug is dark purple and grey. Maps are rendered at 3-4σ. **c,** TWIK2RM R257A:Det (plugged) local resolution showing a central slice through the channel. Select channel elements are labeled. **d,** TWIK2RM R257A:Det (plugged) local B-factor.

**Figure S17** **Mondal, *et al.***

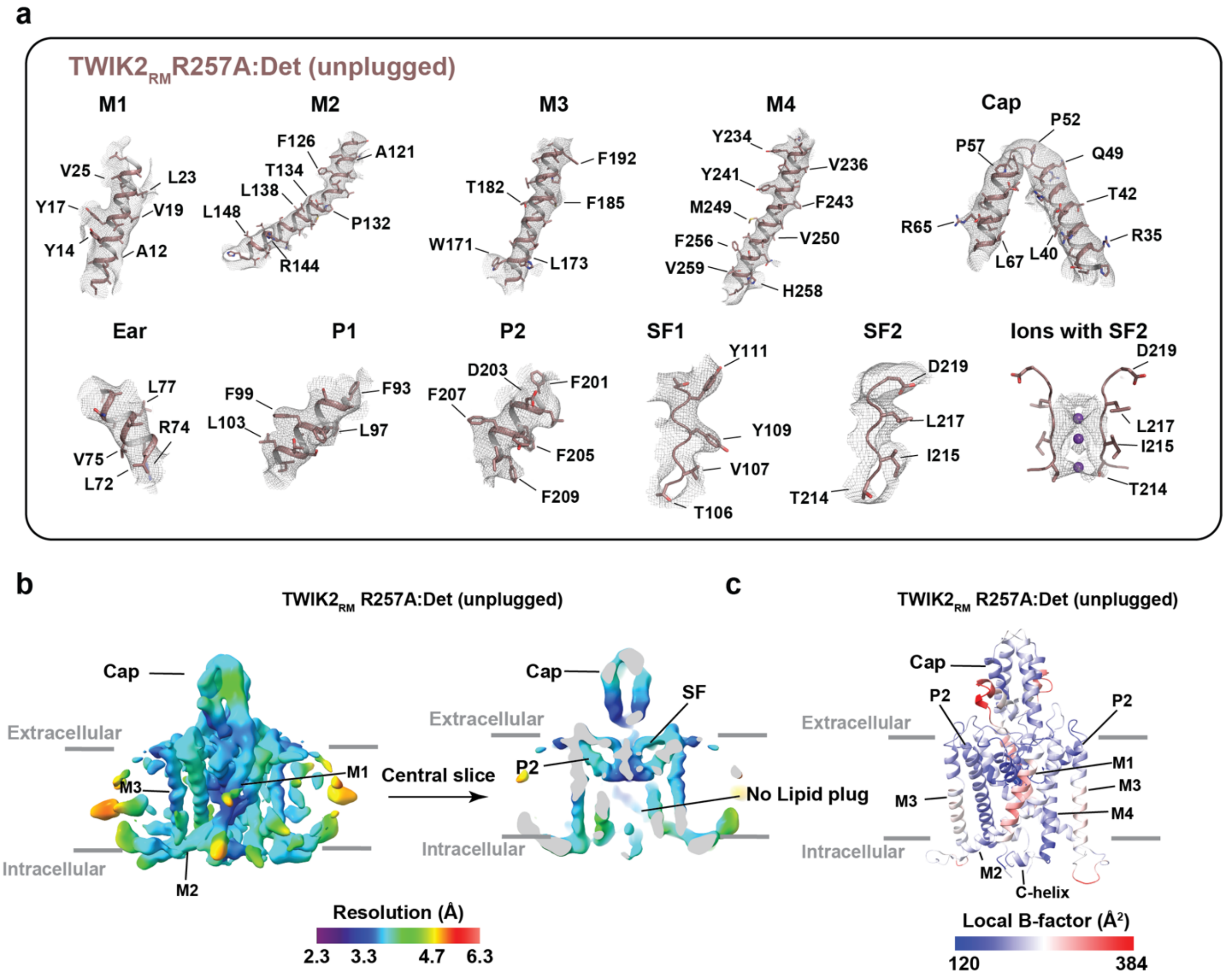


**Figure S17 TWIK2RM R257A:Det (unplugged) cryo-EM map and model quality a**, Electron microscopy maps for indicated TWIK2RM R257A:Det (unplugged) elements. Select residues are indicated. Channel elements are dark purple. Potassium ions are purple. Lipid plug is dark purple and grey. Maps are rendered at 3-4σ. **c,** TWIK2RM R257A:Det (unplugged) local resolution showing a central slice through the channel. Select channel elements are labeled. **d,** TWIK2RM R257A:Det (unplugged) local B-factor.

### Figure S18

Mondal, *et al.*

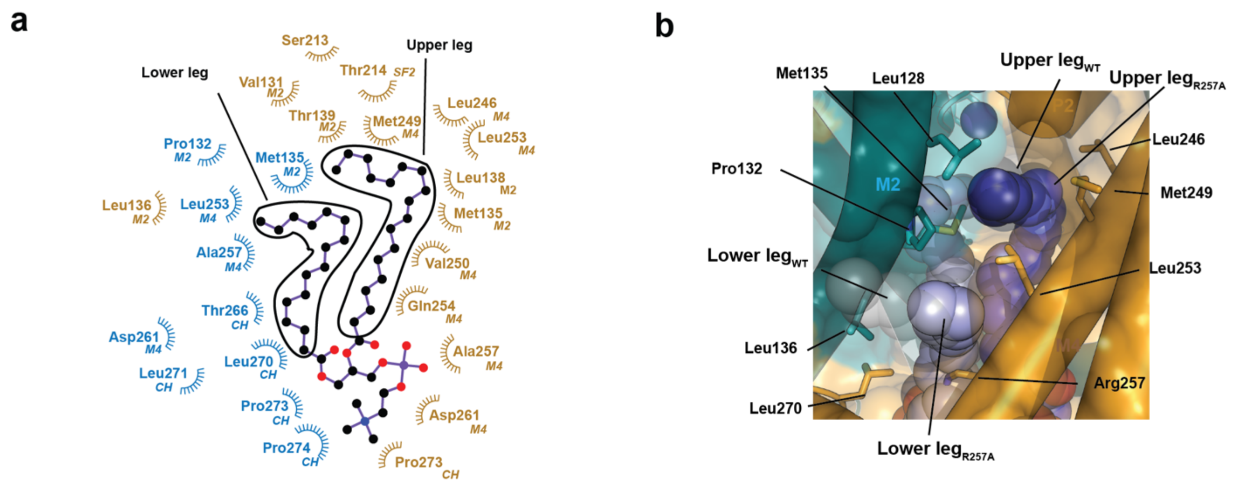


**Figure S18 TWIK2$_{RM}$ R257A :plug lipid interactions. a,** Ligplot[10] diagram showing plug lipid interactions (≤ 5Å). TWIK2$_{RM}$ R257A chains are indicated in blue and olive. Lipid upper and lower leg elements are indicated. Channel elements M2, M4, selectivity filter 2 (SF2), and C-helix (CH) are indicated. **b,** View from the center of the bilayer towards the upper and lower leg binding sites. Comparison of the TWIK2$_{RM}$ R257A plug lipids (slate and light blue) with the TWIK2:ND structure. Lipid plug chains are grey and black and shown in space filling transparent. Interacting residues are shown as sticks.

11 June 2025
